# Cytokinin Senescence Delay Is Shaped by Receptor Specificity and Metabolic Stability

**DOI:** 10.64898/2026.01.12.699116

**Authors:** Omar Hasannin, Risheek R. Khanna, Satyam Singh, Ivan Petřík, Miroslav Strnad, Ondřej Novák, Martin Černý, Aaron M. Rashotte

**Author notes:** Corresponding Author: Aaron M. Rashotte 101 Rouse Life Sciences, Department of Biological Sciences, Auburn University, USA. **Email:** Omar Hasannin, Risheek R. Khanna, Satyam Singh, Ivan Petřík, Miroslav Strnad, Ondřej Novák, Martin Černý, Aaron M. Rashotte.

## Abstract

Each of the four different cytokinin (CK) base forms, *trans* Zeatin (*t*Z), isopentenyladenine (iP), dihydrozeatin (DHZ), and *cis* Zeatin (*c*Z) have distinct chemical metabolism and affinity to the CK Histidine Kinase (CHK) receptors. However, it remains unclear how specific biochemical features of each form such receptor specificity or metabolic differences drives distinct tissue-specific physiological hormone output in response to application of these CK bases. Here, we show that CK receptor preference and metabolic persistence together shape isoform-specific CK signaling strength, including tissue-dependent hormone responses in Arabidopsis leaf versus root assays. Physiological, genetic, and multi-omics integration was used to show that *t*Z and iP anti senescence activity is matched by DHZ through a distinct receptor metabolic mechanism. DHZ requires Arabidopsis Histidine Kinase 3 (AHK3) signaling to be fully effective in a leaf Dark Induced Senescence (DIS) assay and where it overcomes its lower receptor affinity through higher metabolic persistence, accumulating at levels ∼2.5-fold above tZ and iP early in a senescence time course. Together, these findings provide a framework for integration of receptor preference and metabolic stability to determine CK isoform activity.

## Introduction

Plants contain a structurally diverse array of cytokinin (CK) isoforms (Hirose et al. 2008), yet the functional significance of producing and maintaining over 30 different CKs is not fully understood. While general classes of CKs are ascribed with functional roles, such as transport or storage, it is unclear within each class if there are different roles of distinct forms (Sakakibara 2006). Yet, plants invest resources in synthesizing four distinct endogenous base forms, *trans* Zeatin (*t*Z), isopentenyladenine (iP), *cis* Zeatin (*c*Z), and dihydrozeatin (DHZ), alongside massive pools of conjugated base form metabolites (Zürcher and Müller 2016).

The endogenous abundance and distribution of these isoforms vary significantly. In Arabidopsis, active CK bases represent only a small fraction of the total CK pool, approximately 3% of total CKs in shoots and 15% in roots of two-week-old seedlings, with N-glucosides being the most abundant metabolites (Hošek et al. 2020). Among the base forms, *t*Z and iP are the predominant active CKs in Arabidopsis and many dicot species, with endogenous concentrations generally in the range of 0.1–10 pmol g ¹ fresh weight (Novák et al. 2008). DHZ-type metabolites are the least abundant of all CK types in Arabidopsis, approximately 2% of total CKs in shoots (Hošek et al., 2020), and endogenous DHZ base is often below detection limits in standard and organelle-targeted profiling (Antoniadi et al. 2015; Skalický et al. 2023). By contrast, DHZ-type CKs are the predominant free CKs in seeds of legumes such as lucerne and maize (Stirk et al. 2012). Similarly, *c*Z-type CKs are the most abundant CKs in several monocot species including maize and rice, while remaining a minor component in Arabidopsis (Gajdosová et al. 2011).

Active CKs perception and response occur through a well-documented modified Two-Component Signaling (TCS) pathway (Hwang and Sheen 2001; Kieber and Schaller 2018). The prevailing model of CK function remains largely receptor-centric, attributing physiological efficacy of CK forms primarily to ligand-receptor binding affinity (Romanov, Lomin, and Schmülling 2006; Sergey N. Lomin et al. 2015; Spíchal et al. 2004). Under this model, *t*Z and iP are the canonical, “high-activity” CKs due to their high affinity for each of the three ARABIDOPSIS HISTIDINE KINASE (AHK) receptors of CK (Kunikowska et al. 2013; Heyl et al. 2012). Consequently, other base isoforms, DHZ and *c*Z, which have been shown to have lower relative binding affinities are frequently marginalized as “minor” or biologically less relevant. Nonetheless, there have been attempts to identify specific functions of each isoform. Despite lower receptor affinity seen for *cZ* in Arabidopsis and other Eudicot species (Schäfer et al. 2015), it can delay leaf senescence in Monocots such as maize and oat (Behr et al. 2012; Gajdosová et al. 2011). DHZ is even less characterized with reported activity as species and tissue-specific, from less effective activity to being 30-fold more effective in promoting callus growth than *t*Z in some bean species (Mok, Mok, and Armstrong 1978).

Homeostasis is another critical factor affecting the functional activity of CKs which is often less considered when discerning CK activities. Homoeostasis is primarily controlled through irreversible degradation by CYTOKININ OXIDASE/DEHYDROGENASE (CKXs) and also via glucose conjugation through UDP-GLUCOSYLTRANSFERASEs (UGTs) (Werner and Schmülling 2009; Wang et al. 2011). *t*Z and iP are more favorable substrates for CKXs due to the presence of an unsaturated double bond in their N^6^-side chain (Galuszka et al. 2007). In contrast, DHZ, which possesses a saturated side chain, is a significantly less favorable or poor substrate (Galuszka et al. 2007; Reid et al. 2016). This raises questions pertaining to how elimination of DHZ is achieved in plants, which remains to be investigated.

Although these differences are well-known, they are untested as to how these known metabolic distinctions influence physiological outputs. It is unclear whether these minor isoforms merely act as weak mimics of *t*Z or whether interactions between differences in receptor-binding preferences and metabolic stability could drive distinct transcriptional outputs and cause the observed tissue- and species-specific activities for some isoforms, despite sharing the same TCS pathway. To investigate whether receptor specificity and metabolic persistence jointly determine CK isoform activity, we integrated physiological, metabolomic, transcriptomic, and proteomic profiles in leaf senescence as the primary physiological readout. As CK-mediated delay of senescence is one of the most well-characterized CK responses (Gan and Amasino 1995).We demonstrate that physiological efficacy is not dictated by receptor potency alone but by the integration of receptor specificity and metabolic features. We establish that DHZ exhibits robust anti-senescence activity equal to *t*Z, despite its previously determined lower binding affinity. We show this is achieved through a compensatory metabolic strategy, as DHZ metabolic stability allows it to hyper-accumulate, maintaining the concentration threshold required to trigger signaling through an AHK3-dependent pathway.

## Results

### DHZ delays leaf senescence with efficacy comparable to *t*Z and iP

To test the physiological effects of different CK isoforms, we performed dark-induced senescence (DIS) assays on 1µM CK-treated detached Arabidopsis leaves (positions 5 and 6) using Fv/Fm, Chlorophyll (Chl) content, and ion leakage as measures of senescence. The four isoprenoid CK bases *t*Z, iP, DHZ, and *c*Z showed distinct and reproducible differences in their anti-senescence activities **(Fig. 1, Supplemental Fig. S2A, and Supplemental File S1)**. DHZ, *t*Z, and iP consistently demonstrated the strongest anti-senescence activity, while *c*Z showed no statistical difference at any point compared to negative control (NC) **(Fig. 1)**. At 96 hours post-treatment, DHZ-treated leaves maintained an Fv/Fm of 0.72 and a Chl content of 0.83 µmol/mg^-1^, while *t*Z-treated leaves showed an Fv/Fm of 0.68 and a Chl content of 0.70 µmol/mg^-1^ **(Fig. 1A, B)**. In contrast, without CK, NC leaves were reduced to an Fv/Fm of 0.41 and a Chl content of 0.21 µmol/mg^-1^. After 144 hours, *t*Z maintained Fv/Fm of 0.55, and Chl 0.57, while DHZ maintained 0.58 and 0.59 of Fv/Fm and Chl, respectively. Furthermore, in ion leakage measurements at 144 hours, DHZ treatment resulted in the lowest membrane damage (41%), compared to *tZ* (54%), iP (63%), and the NC (80%) **(Fig. 1C)**. To examine the possibility of differential isoform activity in different leaf developmental stages, we repeated the same DIS assay on leaves 3-4 and leaves 7-8 with similar patterns observed across the time course **(Supplemental Fig. S1C-E)**. These results establish a physiological hierarchy in DIS assays of DHZ ≈ *t*Z ≈ iP >> *c*Z ≈ NC. While we noticed a consistent hierarchical trend among base anti-senescence activity in the different measured parameters and leaf positions, DHZ ≥ *t*Z > iP; often, there were no statistical differences between these three isoforms, except in ion leakage, where DHZ was significantly different compared to iP-treated samples.

To determine whether DHZ-activity is concentration-dependent, we performed a dose-response DIS assay. DHZ showed significantly higher Fv/Fm than *t*Z at 1µM and statistically similar levels as *t*Z at 100nM, but was ineffective at 10nM, statistically similar to NC, a concentration at which *t*Z still retained partial activity **(Fig. 1E, F).** This indicates that DHZ high activity is not due to superior potency; its lack of response at 10nM is consistent with its known lower receptor affinity compared to *t*Z. Instead, DHZ requires a high concentration threshold to be effective.

### Unlike tZ, DHZ requires AHK3 to delay leaf senescence

DHZ is generally considered a weak cytokinin because its general affinity for AHK receptors is lower than that of *t*Z and iP (Hluska et al. 2021). Intriguingly, in our DIS assays, DHZ consistently matched their anti-senescence activities (**Fig. 1**). Because it has been previously reported that DHZ preferentially bind to AHK3 over AHK2 and AHK4 (Sergey N. Lomin et al. 2015), we hypothesized that DHZ unexpectedly high activity may depend on signaling through this receptor. We tested this hypothesis using AHK receptor mutants (**Fig. 2A-C**). In *ahk3*, the anti-senescence effect of DHZ was severely reduced (63% and 70% reduction in Fv/Fm and Chl vs WT), compared to *t*Z activity that was relatively less affected (34% and 21% reduction in Fv/Fm and Chl vs WT) **(Fig. 2A-C)**. Conversely, in the *ahk2/4* double mutant, both *t*Z and DHZ activities were reduced to a similar extent, indicating that AHK3 functions as the major factor of DHZ responsiveness.

AHK3 is the only CK receptor known to efficiently phosphorylate ARR2, a key driver of CK-mediated senescence delay (Kim et al. 2006). We therefore tested whether ARR2 is similarly required for DHZ activity. In *arr2*, we found similar trends where DHZ-treated leaves showed 54% and 62% reductions while *tZ*-treated samples were only reduced by 5% and 19% in Fv/Fm and Chl, respectively, compared to WT in DIS.

We found that this AHK3 dependency also explains a tissue-specific activity of DHZ. Unlike DIS assays, in a root inhibition assay, DHZ is a significantly weaker inhibitor of root growth than *tZ* in WT seedlings (*tZ*-treated had 65% shorter primary root than DHZ-treated). However, in *ahk3*, the DHZ inhibitory effect was significantly present, but highly reduced (66% longer than WT), while *tZ* was also less affected (27% longer than WT) **(Fig. 2C and Supplemental Fig. S3)**. This suggests that DHZ is primarily mediated via AHK3, a receptor predominantly expressed in the shoot (Higuchi et al. 2004; Nishimura et al. 2004), making its activity shoot-specific.

### Metabolic persistence compensates for lower receptor affinity in DHZ-treated samples

To connect physiology to CK metabolism, we quantified the endogenous CK levels in treated leaves **(Figure 3; Supplemental File S3)**. All base forms exhibited a gradual decline from 2 to 144 hours post-treatment, consistent with expected metabolic turnover **(Fig. 3A)**. Across treatments, we did not observe substantial interconversion between CK types. Occasional minor accumulations of other CK bases (e.g., < 2 pmol g ¹ FW levels of *tZ* following iP treatment) were detected at isolated time points, but these were small in magnitude and inconsistent over time **(Supplemental Fig. S4 and Supplemental File S3)**.

We found that DHZ maintains the highest concentration throughout the 144-hour time course, 2.5, 1.4, 2.3-fold higher than *t*Z, iP, and *c*Z, respectively, at 2 hours **(Supplemental Table S1 and Fig. 3A)**. To quantify the total exposure and clearance dynamics of each base form in the applied leaf tissues, we fitted a one-phase exponential decay model to the LC-MS time courses. The model provided an excellent fit for *t*Z and DHZ (R² = 0.99, p < 0.01), revealing substantial differences in their persistence under DIS conditions. DHZ displayed a longer apparent half-life (44 h) than *t*Z (23 h), consistent with its known resistance to CKX-mediated degradation. In contrast, the decay model did not significantly fit the *c*Z or iP time courses (R² = 0.72, p = 0.15 and R² = 0.76, p = 0.13, respectively), suggesting that the clearance kinetics of these two bases may not follow simple one-phase decay under our experimental conditions; their half-life estimates (*c*Z: 33 h; iP: 52 h) should therefore be interpreted with caution and are not considered reliable **(Supplemental Table S3 and Supplemental Fig. S5)**. Independent of decay model assumptions, we calculated the cumulative hormone exposure using Area Under Curve (AUC). DHZ provided the largest cumulative exposure, with an AUC approximately 3-fold higher than *t*Z (40,302 vs. 13,847 a.u.) and 2-fold higher than iP (20,273 a.u.). Because DHZ started at 2 hours more than 2-fold higher than the rest of the bases, we normalized the AUC to starting concentration, DHZ retained the highest exposure per unit of initial accumulation (66.5 vs. 44.9 and 47.9 for *t*Z and iP, respectively), indicating that its greater total exposure is not simply a consequence of higher initial uptake but reflects genuinely slower clearance **(Supplemental Table S4)**. In contrast, *t*Z and *c*Z showed similar AUC values (13,847 vs. 14,348 a.u.) despite their different physiological activities, consistent with differences in receptor engagement rather than hormone exposure alone.

This finding is consistent with our dose-response data **(Fig. 1E, F)**, suggesting DHZ prolonged metabolic stability helps it maintain the higher concentrations needed to drive a sustained, AHK3 dependent anti senescence response despite lower affinity. Our regression modeling supports this link. The metabolic profiles of the three bases, *t*Z, iP, and DHZ, were all significant predictors of the anti-senescence response **(Fig. 3B, Supplemental File S4)**. In contrast, the metabolic profile of *c*Z was not a positive predictor. Interestingly, *c*Z was a significantly negative predictor for both Fv/Fm and Chl. Of the non-c*Z* bases, DHZ was the most consistent, being the only form to show significant (p < 0.001) interaction at all time points across all three measured physiological metrics (Chl, Fv/Fm, and Ion Leakage).

As DHZ has been shown to be a less favorable substrate for degradation by CKXs (Reid et al. 2016; Galuszka et al. 2007), this raised the question of how DHZ homeostasis is maintained. We investigated alternative pathways by calculating the rate of conversion of bases to N-glucosides. At 2 hours post-treatment, the amount of DHZ converted into its N-glucoside forms exceeded the remaining detectable DHZ pool at 2 hours, corresponding to 108.2% of the measured DHZ concentration at that time point. A ∼12-fold higher rate than for *t*Z and iP (9.1% and 9.3%, respectively) **(Fig. 3C and Supplemental Table S2)**. This suggests a divergence in metabolic control; while *t*Z and iP are primarily regulated by CKX degradation, DHZ appears to be controlled via rapid N glucosylation.

To genetically support the importance of the clearance through N-glucosylation on DHZ activity, we performed DIS assays on mutants of the UGT CK-N-conjugating enzymes, UGT76C1 and 2. In the ugt76c1/c2 double mutant, where N-glucosylation is >99% blocked (Broke 2021), the anti-senescence effects were massively enhanced. At 144h of DIS, Chl content in DHZ-treated leaves nearly doubled (from 0.49 µmol/mg-^1^ in WT to 0.86 µmol/mg-1 in the mutant). Similarly, for *tZ*, from 0.47 to 0.85 µmol/mg-^1^. In contrast, in the UGT76C2-OE, *t*Z anti-senescence activity was hugely reduced (Chl content 0.19 µmol/mg^-1^), but it still significantly higher than the NC (0.075 µmol/mg^-1^). Meanwhile, DHZ anti-senescence activity was severely reduced to 0.11 µmol/mg^-1^, and was not significant compared to NC. Similar trends were observed for Fv/Fm measurements **(Fig. 3D, F)**. These genetic data suggest that N-glucosylation is important for DHZ homeostasis and activity.

### *t*Z, iP, and DHZ induce overlapping transcriptional programs that suppress senescence

To understand the transcriptional basis for the observed physiological differences, we performed a time-series RNA-seq analysis at 2, 48, 96, and 144 hours post-treatment across DIS examining CK isoforms **(Supplemental File S2)**. Temporal differences were found as the strongest source of variation within the dataset, as seen in a PCA for the results **(Supplemental Fig. S4)**. At 2 hours, *t*Z, iP, and DHZ each triggered a rapid induction of canonical CK-responsive TCS genes, consistent with early CK signaling activation **(Fig. 4B)**. By 48 and 96 hours, these base forms reached their maximal transcriptional impact, as seen by the largest number of Differentially Expressed Genes (DEGs), followed by a sharp decline at 144 hours **(Fig. 3C, D).**

A global comparison of gene expression among base CKs showed that *t*Z, iP, and DHZ induce large and highly overlapping transcriptional changes, while *c*Z is an outlier with minimal regulatory impact on the number of DEGs **(Fig. 3C, D)**. Multi-block Supervised Partial Least Squares Discrimination Analysis (PLS-DA) and PCA further showed that the global expression profiles of the bases *t*Z, and DHZ are more closely similar to each other compared to iP and *c*Z especially at the 48h and 96h time points **(Fig. 3A and Supplemental Fig. S6)**. *t*Z, iP, and DHZ robustly induced core CK-signaling, biosynthesis, and degradation genes **(Fig. 3B)**. Although a slight induction of these genes was observed in cZ-treated samples, this response was generally not statistically significant in our DE analysis **(Fig. 3C, D)**. A similar pattern was observed for CK-golden list genes identified by (Bhargava et al. 2013) where *t*Z, iP, and DHZ elicited comparable induction, whereas *c*Z showed relatively weaker responses, more closely resembling NC **(Supplemental Fig. S5A)**.

To capture global trends in the regulation of processes associated with plant longevity, we averaged z-score expression values for Senescence-Associated Genes (SAGs) and Chlorophyll Catabolic Genes (CCGs) across the treatment time course. *t*Z, iP, and DHZ suppressed the expression of both SAGs and CCGs at 2, 48, and 96 hours before returning to control levels by 144 hours, indicating that transcriptional suppression of these processes precedes the peak window of DE regulation (48-96 hours). *c*Z followed a broadly similar trend; however, very few of these genes reached statistical significance in DE analysis of *c*Z-treated samples. At the individual gene level, expression patterns were largely consistent across *t*Z, iP, and DHZ treatments, except in iP-treated samples, the senescence marker SAG12 exhibited a rebound in expression by 144 hours **(Fig. 3E–G)**.

We next examined the expression of nuclear-encoded chlorophyll biosynthesis genes and plastid-encoded genes. In contrast to SAGs and CCGs, regulation of these gene sets did not begin at 2 hours, expression was comparable across all CK treatments and the NC at this early time point. By 48 hours, however, chlorophyll biosynthesis genes showed a clear upregulation in CK-treated samples relative to NC. Notably, this induction was more pronounced in DHZ-treated samples, particularly at 48 hours. Intriguingly, the expression of plastid-encoded genes showed a divergent response among CKs, in *t*Z- and iP-treated samples, plastid-encoded gene expression was suppressed or remained comparable to NC, whereas in DHZ-treated samples it was significantly elevated at 48 hours, with many of these genes identified as DE relative to NC **(Supplemental Fig. S5 B)**.

### DHZ preferentially sustains photosynthesis-associated gene expression through AHK3

To examine possible isoform-specific regulation, we examined DEGs uniquely regulated by *t*Z, DHZ, and iP. Although the majority of photosynthesis-associated DEGs were shared by all three bases, we found that the unique upregulated DEGs by DHZ were enriched for photosynthesis, light-harvesting complexes, and chloroplast maintenance, whereas *t*Z- and iP-unique DEGs were enriched for general stress-response pathways **(Fig. 4H)**. Across all timepoints we found 305 DEGs shared only between DHZ and *t*Z were enriched for the reductive pentose-phosphate cycle and dark reactions of photosynthesis; notably, RBCS-1A, RBCS-1B, and CFBP1 were included in this set and were not significantly regulated by iP. These distinctions were not apparent at 2 hours but gradually emerged at 48-96 hours, coinciding with the peak transcriptional regulation among treatments.

Overall, we found the average z-score expression of photosynthesis-associated genes, and selected marker genes were higher in DHZ-treated than *t*Z-and iP-treated samples **(Fig. 5A-B)**. To further validate photosynthesis-sustained regulation by DHZ, we performed an unbiased co-expression analysis using only the expression profiles of the three active bases, and correlated the modules with the Fv/Fm, Chl, and ion leakage **(Fig. 5C and Supplemental Fig. S6B-C)**. We identified a gene module (M9) that was strongly correlated with Fv/Fm and Chlorophyll (r > 0.9) and enriched for core photosynthetic components. The expression of this module eigengene was higher in DHZ-treated leaves compared to all other treatments **(Fig. 5D)**.

Furthermore, on an independent set of samples, we used qRT-PCR to validate these findings. In wild-type leaves, DHZ induced the expression of photosynthesis-related genes LHCB1.2 and RBCS (3.0 and 2.9 log2FC) more strongly than *tZ* (2.0 and 2.2 log2FC), while repressing the senescence marker SAG12 to a similar extent (*t*Z: −2.0, DHZ: −1.9 log2FC). In *ahk3*, DHZ ability to regulate these genes compared to WT was significantly reduced (ANOVA post-hoc p < 0.05) for RBCS and LHCB1.2. SAG12 regulation was also reduced to −1.42, but it was not statistically significant (p = 0.267). Whereas *t*Z transcriptional activity remained largely intact, no significant difference between *ahk3* vs WT-col **(Fig. 5E)**. Together, these data support that DHZ strong anti-senescence activity stems from an AHK3-dependent upregulation of photosynthesis-related genes.

In a parallel experiment, we quantified early induction of protein changes by base forms after 2 hours of treatment. We identified 62 Differentially Enriched Proteins (DEPs) across all treatments **(Supplemental File S5 and Supplemental Fig. S9)**. DHZ induced the largest early proteomic response, with 52 DEPs (34 upregulated, 18 downregulated), followed by *t*Z with 21 DEPs (14 upregulated, 7 downregulated). In contrast, iP showed no statistically significant DEPs, and *c*Z produced only 3 **(Supplemental Fig. S10)**. Hierarchical clustering demonstrated that *t*Z- and DHZ-treated samples clustered together and away from iP and *c*Z **(Supplemental Fig. S9)**, indicating that these two bases share a highly similar early proteomic signature.

Of the 52 DHZ DEPs, 14 were shared with *t*Z, and all of those were upregulated in both treatments with complete direction concordance. These shared proteins included several with high fold changes (>3 log2FC), such as AT1G69360, AT3G01920, AT4G22470, and AT4G26390, as well as metabolic enzymes THX, PFK6, SUS4, and LACS8 **(Supplemental Fig. S11)**. For many of these shared proteins, iP and *c*Z showed similar directional trends that did not reach statistical significance.

Among the 38 DHZ-specific DEPs, several were associated with chloroplast function and photosynthesis. These included PORA (protochlorophyllide oxidoreductase A; +1.38 log2FC), which catalyzes a key step in chlorophyll biosynthesis; ndhK (NAD(P)H dehydrogenase subunit K; +0.83 log2FC), a component of chloroplast electron transport; atpF (ATP synthase subunit; +0.81 log2FC); and FTSH1 (+0.60 log2FC), a chloroplast metalloprotease involved in photosystem II repair. *t*Z-treated samples mostly showed similar directional trends for these proteins but did not reach statistical significance **(Supplemental Fig. S11)**. This early enrichment of photosynthesis-related proteins by DHZ is consistent with the later transcriptional enrichment of photosynthetic genes at 48-96 hours and the strong physiological maintenance of Fv/Fm and chlorophyll by DHZ throughout the DIS time course.

## Discussion

Why do plants maintain multiple base forms, such as DHZ and *c*Z, which exhibit lower receptor affinity and are generally considered less active? Here we show that CK isoforms function is not solely dictated by receptor affinity but also relies on the integration of receptor preference and metabolic persistence. We demonstrate that DHZ performs at a level statistically comparable to *t*Z and iP in delaying leaf senescence, establishing a physiological hierarchy of DHZ ≈ *t*Z ≈ iP >> *c*Z ≈ NC. Although the relative ordering DHZ ≥ *t*Z > iP was highly consistent across physiological measures and time points, we did not find statistical difference between the three isoforms **(Fig. 1 and Supplemental Fig. S1)** except for the pairwise comparison of *t*Z versus DHZ at 1 µM that reached statistical significance in the dose-response experiment **(Fig. 1E)**, and Ion leakage (DHZ versus iP). To confirm this observed hierarchy, additional statistical power with more replications would be required to resolve small pairwise differences between these isoforms with high confidence. Nonetheless, DHZ performing at least similar to *t*Z and iP raises questions as to how DHZ, a CK with generally lower receptor affinity (Lomin et al. 2015; Spíchal et al. 2004), can perform on par with high-affinity *t*Z. Our findings here support a two-part, concentration-dependent model as a solution.

The first component of this model is receptor specificity. We showed that DHZ anti-senescence high activity is specialized, dependent on AHK3, the receptor for which DHZ has preferential affinity (Sergey N. Lomin et al. 2015), whereas *t*Z activity is generalized across multiple receptors and assays **(Fig.1 and Fig. 2)**. Aligning with the established role of AHK3 as a primary driver in CK-mediated leaf longevity (Kim et al. 2006). Consistent with this, DHZ anti-senescence activity was severely reduced in *ahk3* mutant, while *t*Z remains largely intact **(Fig 2)**. Furthermore, our transcriptomic and qRT-PCR data show that AHK3 is required for DHZ-preferential and sustained upregulation of specific photosynthesis-related genes **(Fig. 5)**. We also found that DHZ activity is significantly reduced in *arr2* mutant background, consistent with ARR2 functioning downstream of AHK3 in mediating CK responses in senescence (Kim et al. 2006). However, because DHZ activity has not been evaluated in other type-B ARRs mutants, we cannot rule out the possibility that DHZ may also signal through additional response regulators. The contribution of ARR2 relative to other type-B ARRs in DHZ-mediated senescence delay remains to be determined.

This AHK3 dependency also explains the tissue specific activity of DHZ. We found that this high activity by DHZ is not mirrored in root assays, albeit there is still activity in that tissue. This shoot-specific strong activity is likely due to the predominant expression of AHK3 in shoots more than roots (Zhao et al. 2024; Nishimura et al. 2004; Higuchi et al. 2004). These results align with previous literature that DHZ is capable of activating AHK3 and its orthologs in other species (Hluska, Hlusková, and Emery 2021; Vinciarelli et al. 2025). But the implication of this AHK3 specificity has not been shown before. *t*Z activity remains functional in the *ahk3* background. Demonstrating that *t*Z can effectively delay senescence by signaling through AHK2 and AHK4 as has been shown in previous reports (Riefler et al. 2006).

This AHK3-dependency, however, does not by itself, explain DHZ high activity, as *t*Z is known to have an even higher affinity for AHK3. We found that DHZ activity is not driven by potency, but by metabolic persistence and increased total exposure compared to *t*Z and iP **(Fig. 4)**. In dose response DIS assays, at high concentrations (1µM and 100nM), DHZ anti-senescence activity is robust. But at a low, threshold concentration (10nM), its effect is lacking, while high-affinity *t*Z retains partial function **(Fig. 1E)**. Suggesting that DHZ activity requires higher concentration to overcome its lower receptor affinity. CK metabolic profiling recapitulates this finding, while *t*Z is rapidly degraded by CKXs (Galuszka et al. 2007; Reid et al. 2016), DHZ is highly resistant and persists at concentrations 2.5-fold higher than *t*Z after 2 hours of treatment **(Fig. 3A and Table S1)**. Together, these findings provide an explanation for why DHZ, in certain tissues and species, possess higher or similar activity to *t*Z (Mok et al. 1978).

Although iP displayed higher total exposure compared tZ (**Fig. 3A**), it remained relatively less effective in delaying DIS than *t*Z. This indicates that metabolic persistence is necessary but not sufficient for strong anti-senescence activity. Under a model where AHK3 is a major senescence-related receptor, receptor specificity provides a plausible explanation that iP has lower affinity to AHK3 than either *t*Z or DHZ. Thus, despite its higher total exposure, iP fails to reach the effective receptor occupancy in AHK3-dependent tissues that *t*Z and DHZ achieve. Interestingly, the clearance kinetics of both iP and cZ were not well described by a simple one-phase decay model, unlike tZ and DHZ which fit the model well (R² = 0.99). This may reflect additional biological complexity in the turnover of these isoforms, such as interconversion, subcellular compartmentalization, or transport, processes that would not be captured by a single exponential model and that warrant further investigation.

As an alternative pathway to degradation by CKXs, we show that DHZ goes through rapid N-glucosylation at an astonishing 11.8-fold higher rate than *t*Z (108.2% vs. 9.1%) **(Fig. 3C).** This metabolic divergence is supported by our genetic data **(Fig. 3D)**. When this N-glucosylation pathway is blocked (in the *ugt*-dm), DHZ anti-senescence activity is enhanced. More importantly, our UGT76C2-OE data genetically recapitulates the 10nM dose-response experiment. By biologically driving the free DHZ concentration below its functional threshold, the UGT-OE line causes DHZ activity to reduce to a level similar to NC. In contrast, high-affinity *t*Z, while reduced, maintains partial function, which was statistically significant compared to NC **(Fig. 3D)**. This suggests that the sustained high concentration allows DHZ to trigger AHK3-pathway, compensating for its poor affinity.

Although this study was conducted in *Arabidopsis thaliana*, the proposed framework is likely applicable across species. However, because both receptor affinities and isoform abundances differ between species (S. N. Lomin et al. 2012), the framework would predict different activity in different plants rather than a universal ranking of isoforms. In Arabidopsis as we have shown, DHZ compensates for lower receptor affinity through metabolic persistence to achieve high anti-senescence activity via AHK3. In other species, where metabolic features and binding preferences differ, a different isoform may occupy that role.

Importantly, it remains to be tested how these dynamics play out endogenously. Our experiments relied on exogenous application of CKs. More broadly, the model raises the possibility that plants may deploy metabolically stable isoforms like DHZ for processes requiring sustained CK signaling, such as prolonged maintenance of photosynthetic capacity during grain filling (Chen et al. 2020). While reserving high-potency, rapidly cleared isoforms like *t*Z for acute, transient responses. Previous targeted studies have shown endogenous changes in isoform concentrations, such as (Matsuo et al. 2012) where DHZ have shown more dynamic changes in concentrations relative to other measured isoforms in tomato ovaries, albeit, the concentrations were much lower than the detected *t*Z. A more systematic quantification of isoform-specific CK dynamics across developmental stages and tissues would be needed to test this hypothesis.

In summary, our findings establish that CK isoform activity is determined by the integration of receptor specificity and metabolic persistence rather than receptor affinity alone **(Fig. 6)**. This framework explains how DHZ which traditionally has been classified as a minor or weak cytokinin, achieves anti-senescence activity comparable to the canonical high-affinity forms *t*Z and iP. DHZ accomplishes this through a compensatory strategy: preferential signaling through AHK3, the key senescence-related receptor, combined with metabolic resistance to degradation that sustains the hormone concentrations needed to drive signaling despite lower binding affinity. More broadly, this model provides a basis for predicting isoform-specific CK activity across tissues and potentially across species, moving beyond simple affinity rankings toward an integrated view where the biochemical properties of each isoform contribute to its activity.

## Materials and Methods

### Planting material and growth conditions

Wild-type Arabidopsis seeds (Col-0) were planted in plastic trays with Pro-Mix BX and placed in growth chambers under a 16/8h light cycle (100 μmol m^−2^ s^−1^) with temperatures of 22°C/18°C. *ahk3*, *arr2* were seeds obtained from the ABRC stock center (CS2103396, CS6974), *ugt76c1,c2* double mutant and UGT76C2 OE lines were a generous gift from Dr. Tomas Werner at Graz University in Austria (Brock 2021), *ahk2,4* double mutant line from Jan Hejátko at CETITEC in the Czech Republic.

### Dark-induced leaf senescence and root inhibition assays

Dark-induced leaf senescence (DIS) bioassays were modified for use with Arabidopsis as previously described in (Fletcher and McCullagh 1971; Hallmark et al. 2020; Johnston et al. 2026). Briefly, 4-week-old leaves were detached and floated on 3 mL of 3 mM MES buffer solution (pH 5.7) in 6-well culture plates, one leaf/well; only positions 5 and 6 were used unless otherwise indicated. Cytokinin treatments (tZ, iP, DHZ, and cZ) were applied individually to a final concentration of 1 µM and compared to DMSO or Ethanol (0.1%) as a vehicle control and placed under growth chamber conditions in the dark. Measurements of DIS were performed at 2, 48, 96, or 144 hours after treatment/dark incubation, as noted below. All CKs were obtained from OlChemIm (Olomouc, Czech Republic) as analytical standards with >95 % purity and with N-glucosides free from base form contamination. Adenine was obtained from AmBeed Co. with >99% purity.

For root inhibition assays, WT-Col seeds were sterilized using 10 min ETOH 70% incubation and 20% bleach and tween for 10 minutes followed by sowing on 1x MS and + 1% sucrose pH 5.7 medium, then stratification at 4°C for 2 days. Seedlings were then grown for 4 days under (16/8h) light cycle and transferred to MS supplemented with a final concentration of 1µM of the indicated treatments, and primary root tip was marked, and then primary root growth measurements were taken after 5 days of incubation in the supplemented medium. Measurements were taken from at least 14 seedlings for each replicate/condition, repeated 3 times for each condition; the total number of seedlings measured, n ≥ 38/condition. ANOVA followed by post-hoc was used to determine statistical differences.

### Photosystem II efficiency (Fv/Fm), chlorophyll content ion leakage measurements

In a dark room, FluorCam FC 1000-H was used to measure Fv/Fm; leaf areas were selected manually, and the scaling bar was set to 100. Chlorophyll (Chl) was extracted according to (Nayek et al. 2014; Johnston et al. 2026). Briefly, leaf weights were recorded at the start of the experiment. Leaves were harvested at each of the designated time points, followed by Chl extraction using methanol incubation overnight. Debris was pelleted by centrifugation at maximum speed for 10 min, and supernatant was used to measure total Chl normalized to fresh weight. For ion leakage measurement, detached leaves were washed three times with deionized water, followed by immersion in deionized water, then gently shaken for 2 h at room temperature. Total conductivity was measured as initial readings data, then samples were boiled and cooled down to room temperature and measured again with a bench-top conductivity meter. Total electrolyte leakage is determined by the following formula: Ion leakage (%) = initial/final conductivity 100 (Zhang and Guo 2018). The number of biological replicates is indicated in the figure legend of each experiment; generally, at least four independent biological replicates were performed per treatment/time-point. ANOVA followed by post-hoc test was used to determine statistical differences.

### Quantification of cytokinin profiles

Whole leaf samples (20-30 mg fresh weight) were ground in liquid nitrogen then lyophilized and stored until quantification was performed using three biological replicates (n=18 independent leaves per replicate). Lyophilized tissues were extracted in 0.5mL extraction solvent consisting of 5% formic acid in 75% methanol (both v/v). Four zirconium oxide homogenization beads and a mixture of stable isotope-labelled internal standards of CKs were added to each sample (0.2 pmol CK bases, ribosides and N-glucosides, 0.5 pmol CK O-glucosides and nucleotides). The standards were obtained from Olchemim Ltd. (Olomouc, Czech Republic) and from the Faculty of Science, Palacký University in Olomouc (Czech Republic) with distinct compounds as listed in (Svačinová et al. 2012). The samples were extracted using mix-mode solid phase extraction (Dobrev and Kamínek 2002). First, the samples were shaken in Retsch MM 400 oscillation bead mill (Retsch, Haan, Germany) for 5 min at 27 Hz, 4°C, sonicated for 3 min and incubated for 30 min at 4°C. Then the extracts were centrifuged at 20,000 rpm, 4°C for 15 min (Allegra 64R benchtop centrifuge, Beckman Coulter, USA). The supernatant was diluted with 2.5 mL 1 M aqueous formic acid and loaded onto activated Oasis® MCX 30 mg/1 cc extraction cartridge (Waters, Milford, USA). The activation was performed using 1 mL methanol, and 2mL 1 M aqueous formic acid. After the loading of the sample, the cartridge was washed with 1 mL 1M aqueous formic acid and 1 mL 80% methanol (v/v). The CKs were eluted with addition of 1mL aqueous 0.35 M ammonia and 2 mL 0.35 M ammonia in 60% methanol (v/v). The eluate was evaporated to dryness using SpeedVac concentrator (RC1010 Centrivap Jouan, ThermoFisher, USA) and reconstituted in 40 µl of 10% aqueous methanol (v/v). The sample was transferred into LC vial equipped with a glass insert (Chromservis Ltd., Czech Republic) and then analyzed using ultra-high performance liquid chromatography Acquity I-Class system (Waters, Milford, USA) coupled with Xevo TQ-S series tandem mass spectrometer (Waters, Manchester, UK). The chromatographic conditions and mass spectrometry settings were set as previously published (Svačinová et al. 2012). Data were acquired and processed in multiple reaction monitoring using MassLynx V4.2 software. The final concentration of plant hormones (pmol/g of fresh weight) was calculated using isotope dilution method (Rittenberg and Foster 1940). Values were then log-transformed, and non-detected values were replaced with the detection limits to enable statistical comparisons determined by ANOVA followed by a post-hoc test.

### RNA extraction Transcriptome analysis

RNA was isolated from frozen whole leaf powder (approximately 60 mg) of three biological replicates, n=18 leaves per 1 biological replicate; three independent biological replicates were extracted using RNeasy Plant Mini Kits (Qiagen) according to the manufacturer’s instructions. RNA was sent to Novogene, Inc for Illumina sequencing.

Raw RNA-sequencing reads were processed by Novogene, including raw gene counts, differential expression of genes between treatments, and GO enrichment. Briefly, high-quality clean transcript reads for all downstream analyses were obtained by first removing sequencing adapters, reads containing poly-N, and low-quality reads. Using HiSAT2 *v2.0.5* (Pertea et al. 2016), the *Arabidopsis thaliana* TAIR10 reference genome was indexed and paired-end clean reads were aligned. The mapped reads were assembled by StringTie *v1.3.3b* in a reference-based approach and counted using featureCounts *v1.5.0-p3* (Liao, Smyth, and Shi 2014; Pertea et al. 2015). Differential expression analysis of two conditions/groups (three biological replicates per condition) was performed using the DESeq2 R package (1.20.0) (Love, Huber, and Anders 2014). The resulting P-values were adjusted using the Benjamini and Hochberg’s approach for controlling the false discovery rate. Genes with an adjusted P-value (padj) ≤0.05 found by DESeq2 were assigned as differentially expressed (Robinson, McCarthy, and Smyth 2010). The P values were adjusted using the Benjamini & Hochberg method and a corrected P-value of 0.05 and absolute fold change of 2 were set as the threshold for significantly differential expression.

### Regression Analysis for the integration of gene expression, metabolic data, and physiology

High-variance genes were identified by calculating the variance of mean logFPKM values across all experimental conditions. We first filtered to retain genes in the top 33.3% of variance (quantile > 0.667). From this subset, the top 10,000 genes with the highest variance were selected for further analysis. To identify gene expression signatures that discriminate between cytokinin treatments and controls across the experimental time-course, we used a multiblock Partial Least Squares Discriminant Analysis (block.splsda) using the mixOmics package (Rohart et al. 2017). Expression data (z-scores) were split into four discrete blocks corresponding to sampling time points (2, 48, 96, and 144 h). A full weighted design matrix was implemented to maximize correlations between temporal blocks. Prior to modeling, features with near-zero variance were removed.

Co-expression analysis was conducted using the R package tidyverse, according to (Li, Deans, and Buell 2023). Nodes and Edges selection were calculated using Empirical determination using rank distribution. Edges with r>0.9 and FDR less than 0.0001 were used in the analysis. Only modules containing more than 5 genes were retained. To correlate gene expression with physiological responses, in **(Fig. 5E and Supplemental Fig. 5A, B)**, we conducted OPLS using ropls (version 1.39.0) R package to correlate modules with physiological responses. Modules were selected based on significant correlations with physiological traits. Module 9, which is enriched for photosynthesis-related genes (323 genes), showed a strong positive correlation with Fv/Fm (R = 0.92) and chlorophyll content (R = 0.87), and a strong negative correlation with ion leakage (R = −0.90; all p < 0.0001) **(Supplemental File S4)**.

To analyze the effects of the metabolic profiles of cytokinin forms and their interactions with timepoints on Fv/Fm, chlorophyll content, ion leakage, and gene expression, we used a Generalized Linear Mixed Model (GLMM). The analysis was performed using the glmer function from the lme4 package in R (version 1.1-34). Prior to the analysis, all continuous predictors were scaled (mean-centered and standardized to unit variance). Cytokinin concentrations were fixed effects (predictors) accounted for interactions with time points, while treatments were included as a random intercept to account for variability across experimental treatments. A Gamma distribution with a log-link function was chosen to model the data due to the skewed distribution of chlorophyll content, Fv/Fm, and Ion leakage. Model optimization was carried out using the bobyqa optimizer, with the maximum number of function evaluations set to 100,000 to ensure convergence. Model fit and performance were evaluated using Akaike Information Criterion (AIC) and Bayesian Information Criterion (BIC) to compare models and assess relative goodness of fit. Additionally, variance inflation factors (VIFs) were calculated to check for multicollinearity, and overall model performance metrics were assessed using the model_performance function from the performance package in R. Coefficients were exponentiated for interpretability **(Supplemental File S4)**.

### Quantitative RT-PCR

Treated detached leaves (positions 5-6) were incubated in the dark for 96 hours for (**Fig. 5E**) At least 2 leaves per treatment/condition were ground together and counted as one biological replicate (repeated 3 times). Followed by RNA extraction as previously described. 500 ng of RNA was used for cDNA synthesis using Quantbio qScript cDNA Synthesis Kit. 5 µL of 1:10 dilution of cDNA was used for qPCR mix, using PerfeCTa SYBR® Green FastMix. All primers used in the study are in **(Supplemental File S6)**. TUB4 served as an internal control; relative expression was calculated using ΔΔCt method. Statistical analysis was conducted by comparing ΔCt values using ANOVA post-hoc.

### Proteome analysis

The extraction of proteins from lyophilized tissues was conducted according to the protocol delineated by (Berková et al. 2023). In brief, tissue samples underwent homogenization, and aliquots ranging between 10-20 mg were lyophilized. Subsequently, these were extracted utilizing a mixture of tert-butyl methyl ether and methanol (3:1 ratio). The precipitated protein pellets were then resolubilized in a solution composed of 8 M urea, 10 mM dithiothreitol (DTT), and 100 mM ammonium bicarbonate. Following solubilization, the proteins underwent alkylation and were digested with trypsin. Each sample, containing precisely 5 µg of peptides, was subjected to analysis via nanoflow reverse-phase liquid chromatography coupled with mass spectrometry (LC-MS/MS). The chromatographic separation was achieved using a 15 cm C18 Zorbax column on an Agilent system, interfaced with a Dionex Ultimate 3000 RSLC nano-UPLC and an Orbitrap Fusion Lumos Tribrid Mass Spectrometer, equipped with a FAIMS Pro Interface (Thermo Fisher Scientific, Waltham, MA, USA). The analysis utilized alternating FAIMS compensation voltages of −40, −50, and −75 V. The acquired MS/MS spectra were subsequently recalibrated and interrogated against the Araport 11 protein database, alongside a database of common contaminants, using Proteome Discoverer 2.5. Algorithms such as Sequest HT, MS Amanda 2.0, and MSFragger facilitated the search (Dorfer et al. 2014; Kong et al. 2017). Quantitative analysis was restricted to proteins identified by at least two unique peptides The comprehensive mass spectrometry proteomics dataset has been made accessible through the ProteomeXchange Consortium via the PRIDE partner repository, under the dataset identifier PXD053034. Differential Enriched Protein analysis was conducted using DEP package in R.

## Author contributions

O.H. and A.M.R. designed the experiments. O.H. and R.R.K. collected and prepared samples for RNA sequencing, proteomic analysis, and cytokinin metabolic profiling. O.H. conducted physiological experiments and integration of the omics dataset. S.S. conducted root assay. M.C. performed and analyzed LC-MS proteomic data. I.P., M.S., and O.N. conducted cytokinin metabolic measurements. O.H. and A.M.R. wrote the manuscript, with input from co-authors.

## Supplementary Data

Supplemental Figure S1. Quantification of senescence progression in different leaf developmental stages.

Supplemental Figure S2. Representative images of detached leaves from 4-week-old Arabidopsis plants positions 3-4 and 7-8 treated with 1 µM of cytokinin bases across DIS.

Supplemental Figure S3. Representative images of root length assay of seedlings treated with 1 µM tZ or DHZ for 5 days.

Supplemental Figure S4. Correlation matrix of endogenous cytokinin metabolites across all treatments and time points.

Supplemental Figure S5. Exponential decay model of base form concentrations over the DIS time course.

Supplemental Figure S6. Principle Component Analysis of base form transcriptional profiles across time.

Supplemental Figure S7. Expression profiles of cytokinin-responsive and photosynthesis-related gene sets in base-treated samples.

Supplemental Figure S8. OPLS analysis correlating gene co-expression modules with physiological parameters.

Supplemental Figure S9. Clustering analysis of early proteomic profiles of CK-treated samples.

Supplemental Figure S10. Volcano plots of differentially accumulated proteins in base form–treated samples compared to negative control.

Supplemental Figure S11. Protein expression profiles of shared and DHZ-specific differentially enriched proteins across CK treatments.

Supplemental Table S1. Median base accumulation in base-treated samples across time

Supplemental Table S2. Conversion rate of base forms into N-glucosides

Supplemental Table S3. One-phase exponential decay model fitted to LC-MS time courses of applied CK bases during DIS

Supplemental Table S4. Cumulative hormone exposure of applied CK bases over the DIS time course

Supplemental File S1. Physiological measurements.

Supplemental File S2. Raw gene counts, Differentially Expressed Genes, shared and unique DEGs for active forms.

Supplemental File S3. Cytokinin profiling across the time course of treated samples. Supplemental File S4. Co-expression modules and regression analysis

Supplemental File S5. Proteome data, differentially abundant proteins and raw protein abundances

Supplemental File S6. qPCRs, and primers

## Data availability

All data are incorporated into this article and its online supplementary material. Customized R scripts are in the GitHub Repository (https://github.com/OmarHas07/RNA-ProteomeAnalaysis)

---

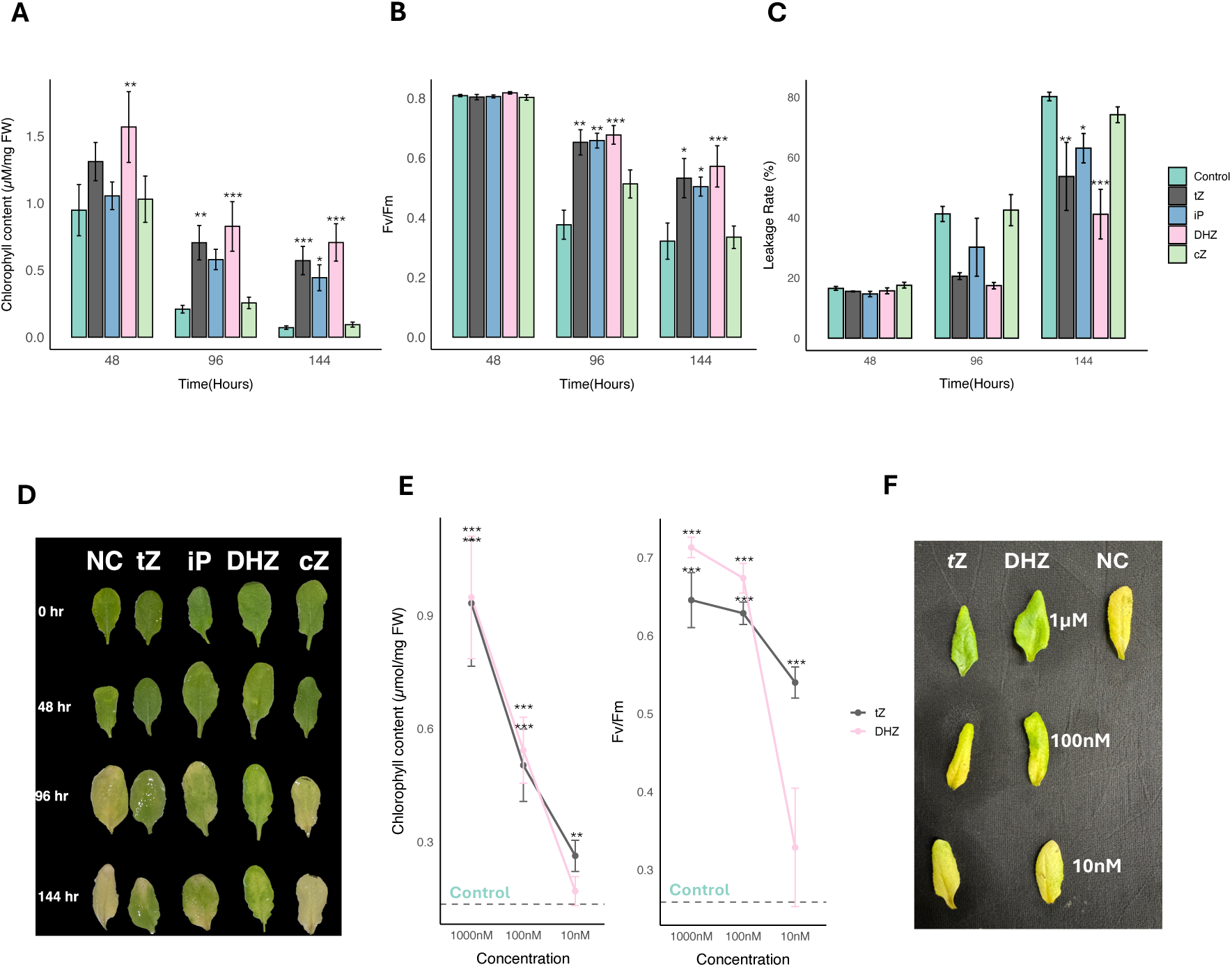

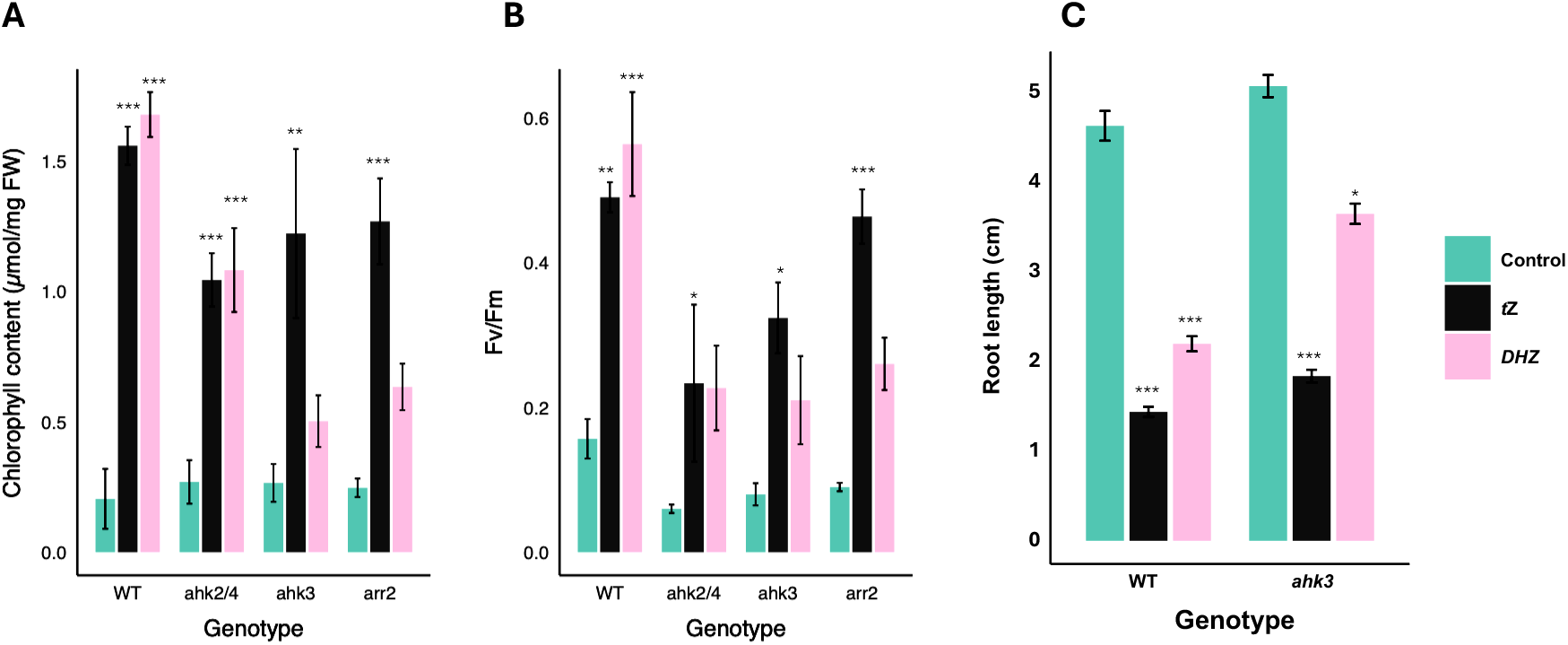

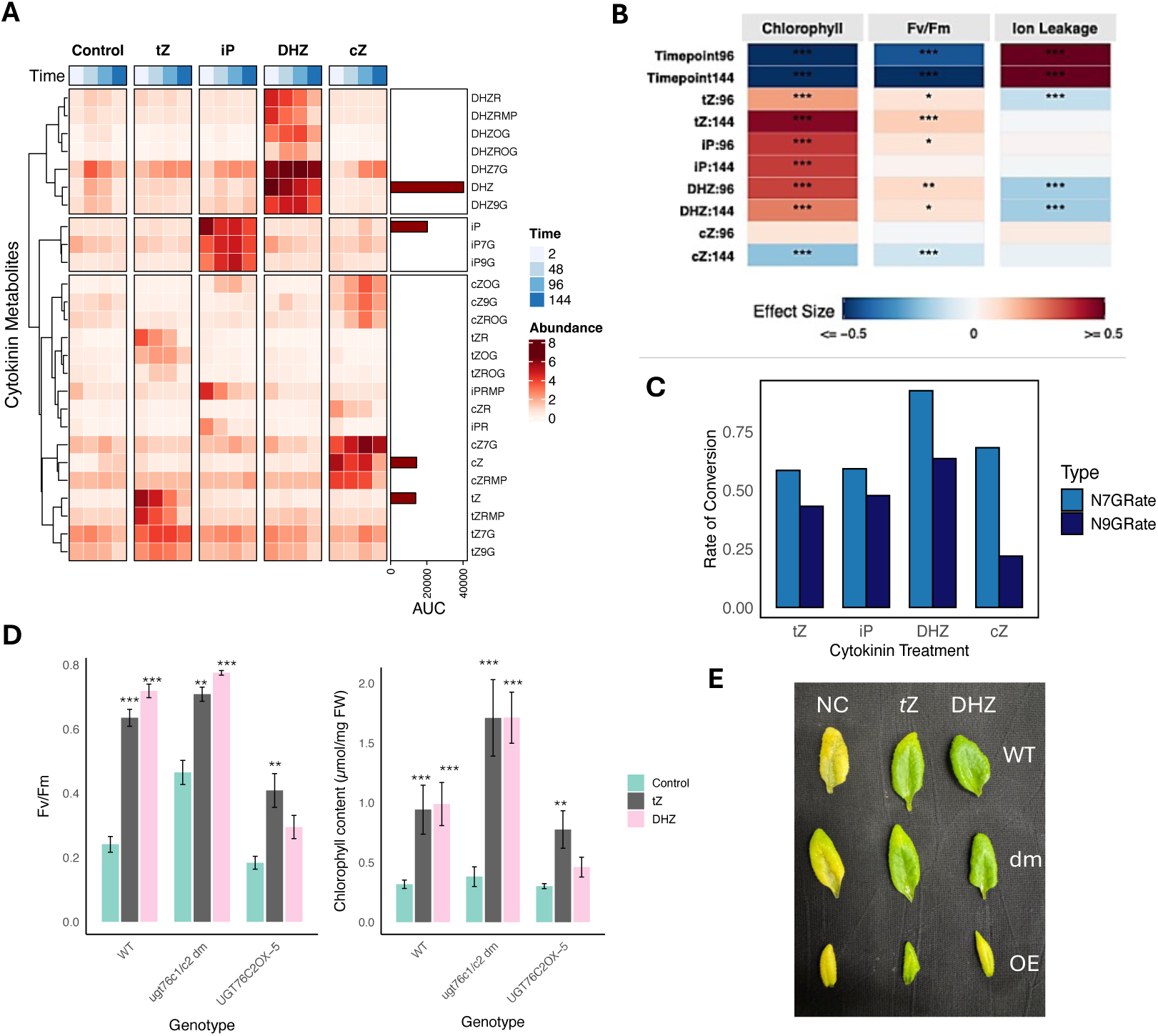

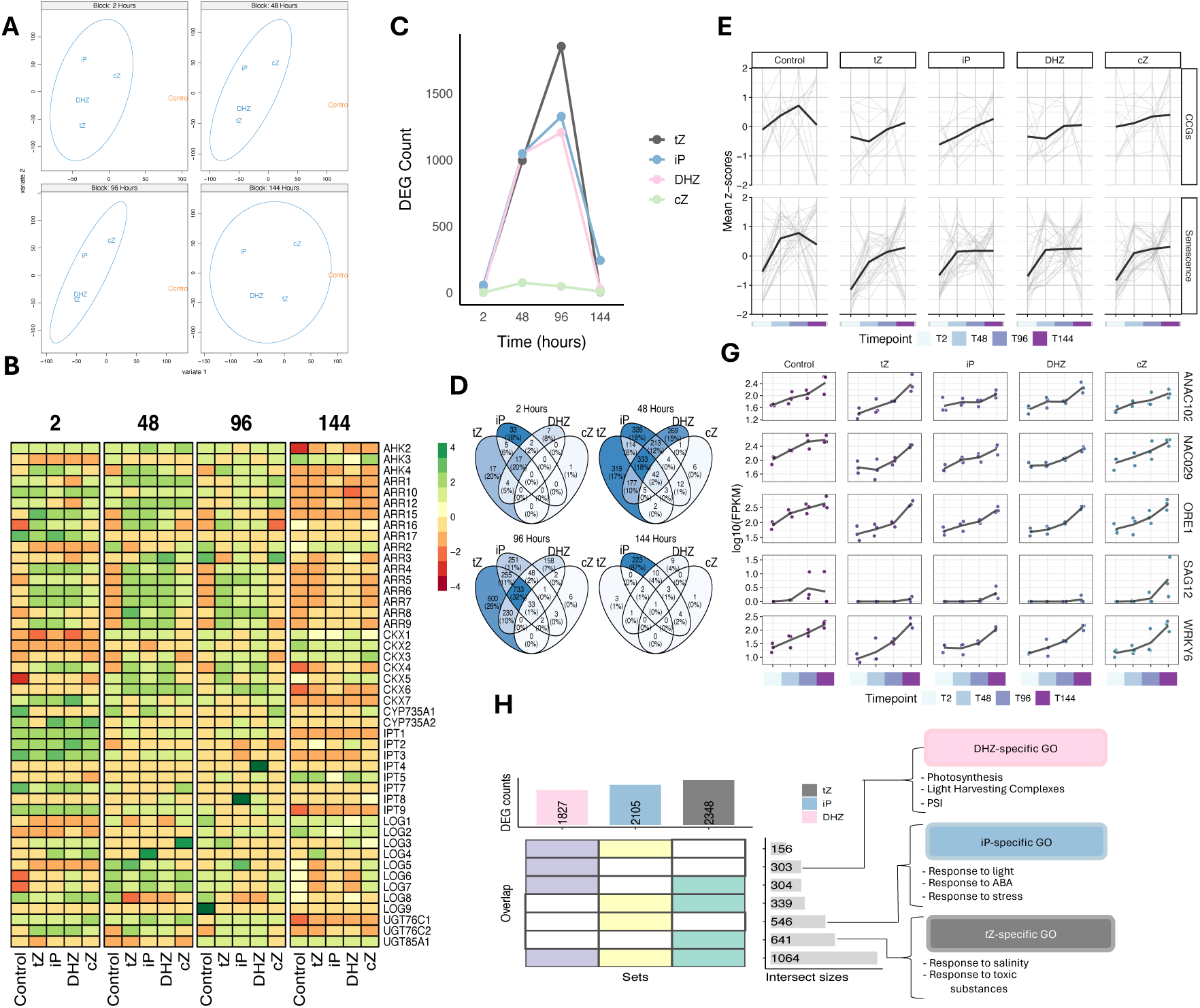

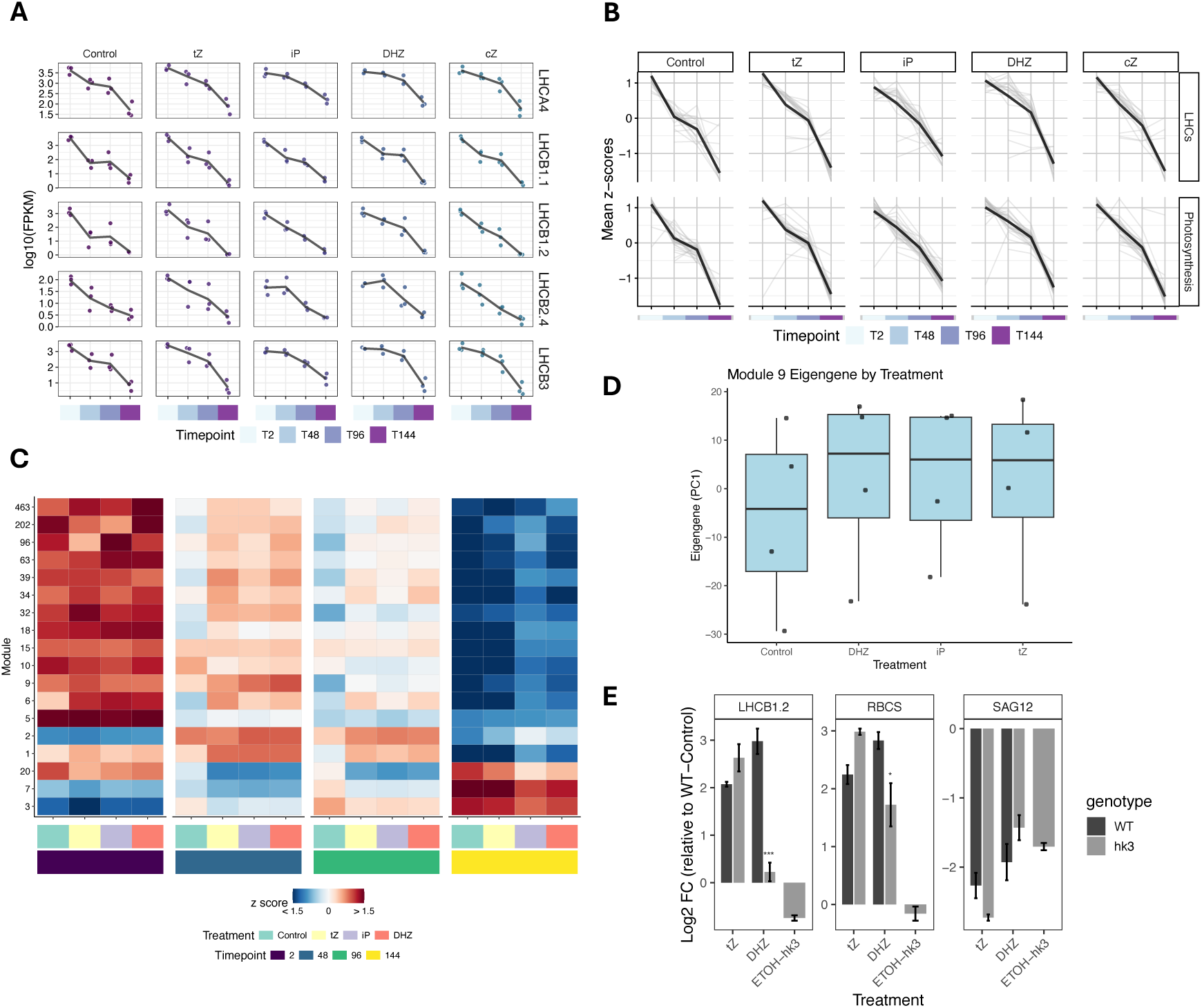

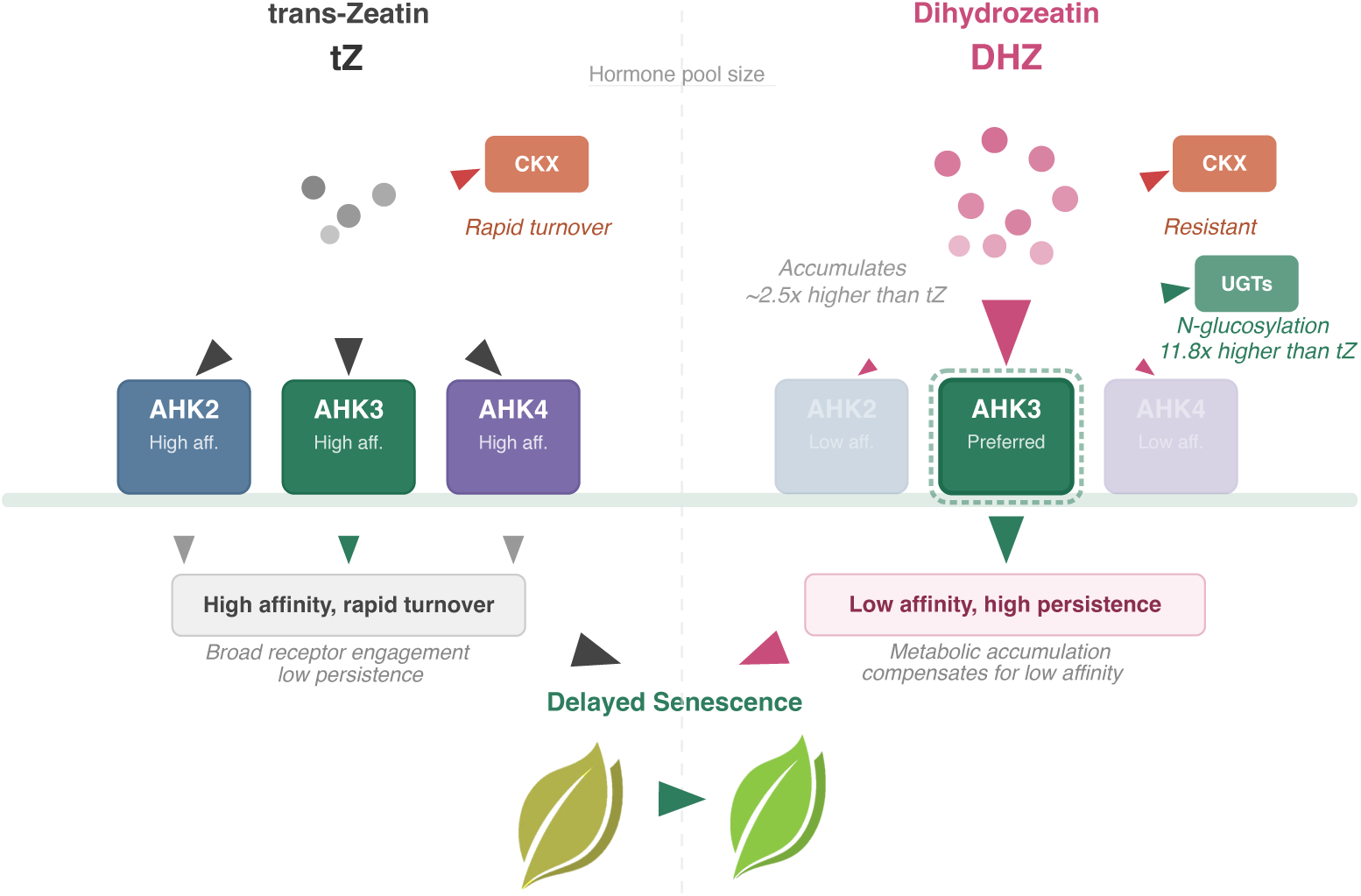

