## Supplemental Figures for "Cytokinin Senescence Delay Is Shaped by Receptor Specificity and Metabolic Stability"

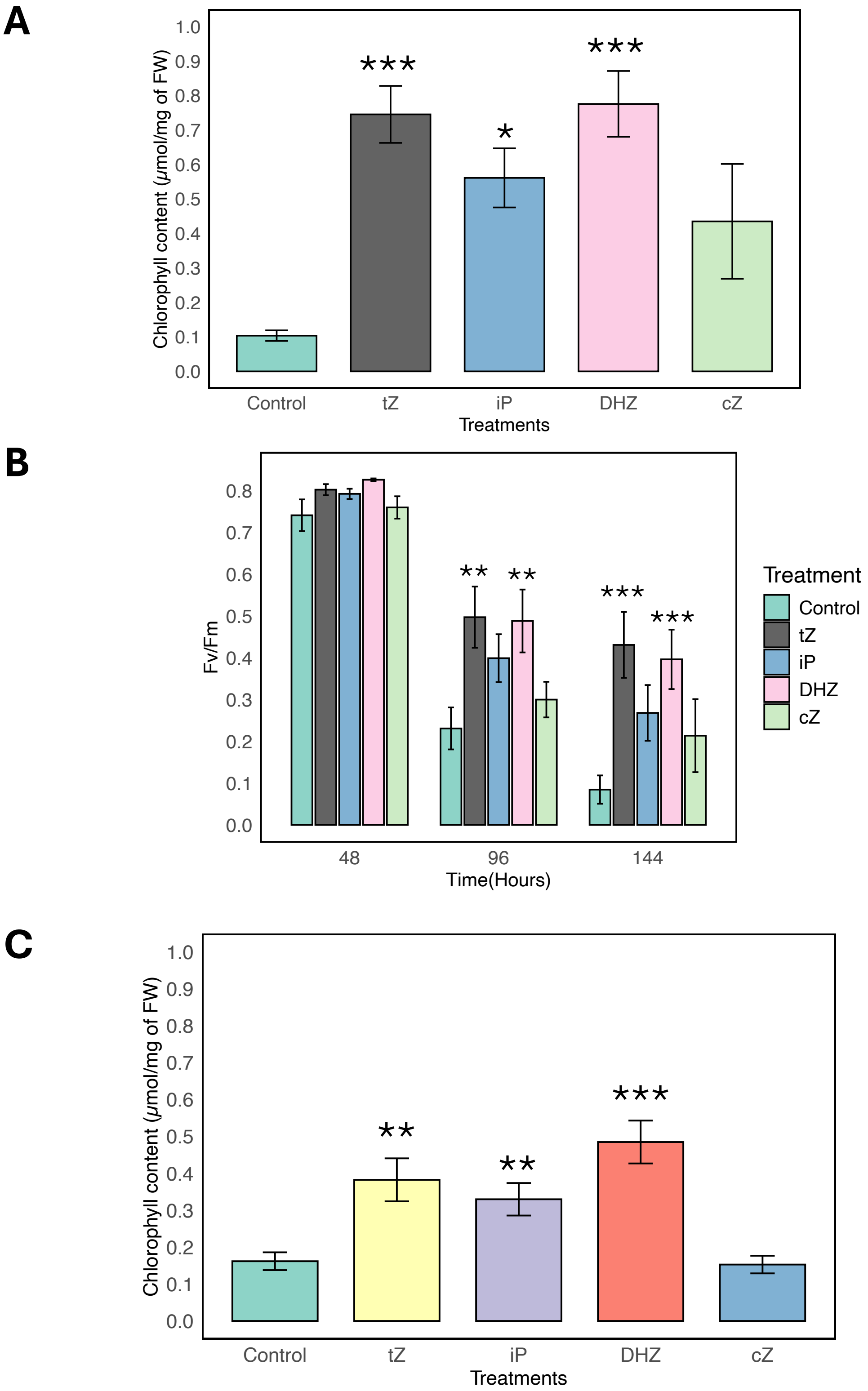
A
Chlorophyll content (μmol/mg of FW)
1.0
0.9
0.8
0.7
0.6
0.5
0.4
0.3
0.2
0.1
0.0
***
*
***
Control
tZ
iP
DHZ
cZ
Treatments
B
Fv/Fm
0.8
0.7
0.6
0.5
0.4
0.3
0.2
0.1
0.0
**
**
***
***
48
96
144
Time(Hours)
Treatment
Control
tZ
iP
DHZ
cZ
C
Chlorophyll content (μmol/mg of FW)
1.0
0.9
0.8
0.7
0.6
0.5
0.4
0.3
0.2
0.1
0.0
**
**
***
Control
tZ
iP
DHZ
cZ
Treatments

**A**

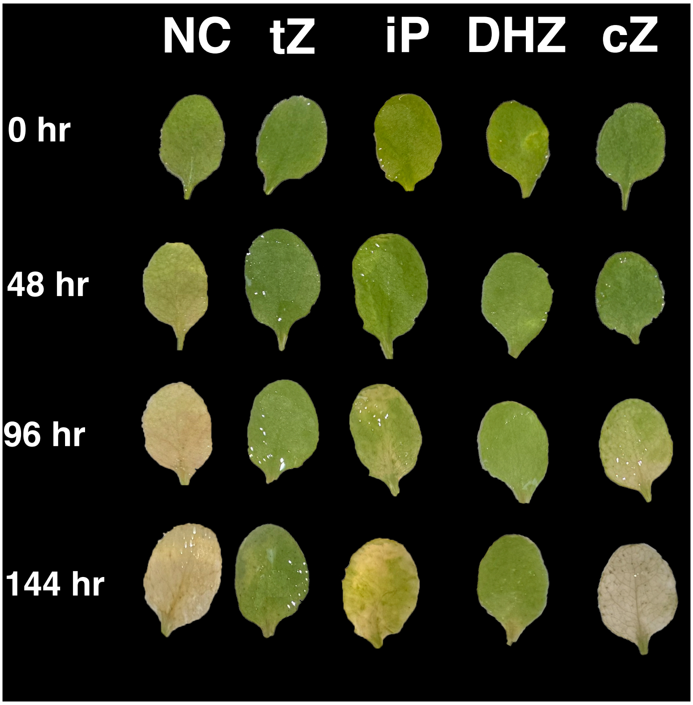


**B**

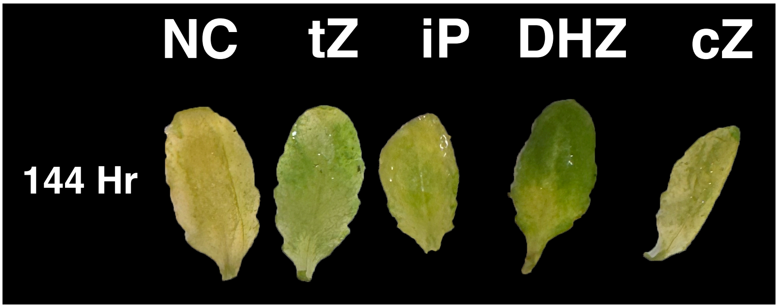

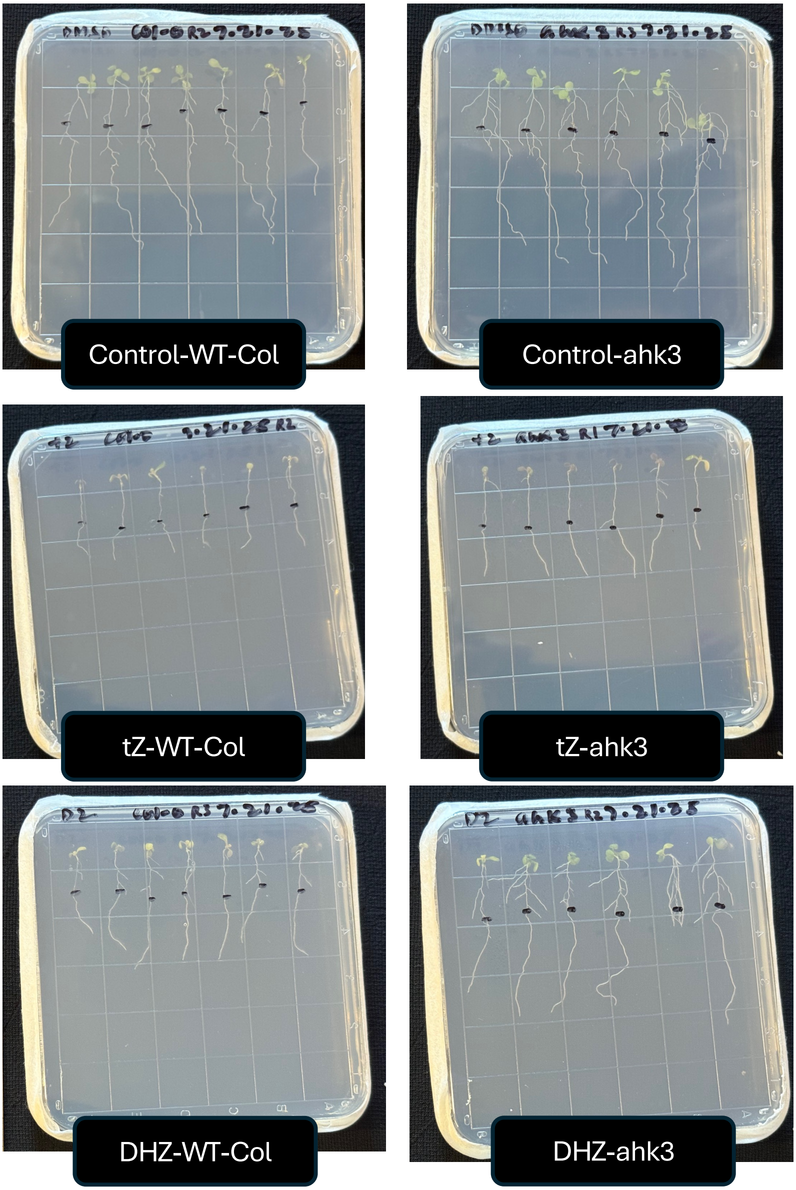
Control-WT-Col
Control-ahk3
tZ-WT-Col
tZ-ahk3
DHZ-WT-Col
DHZ-ahk3

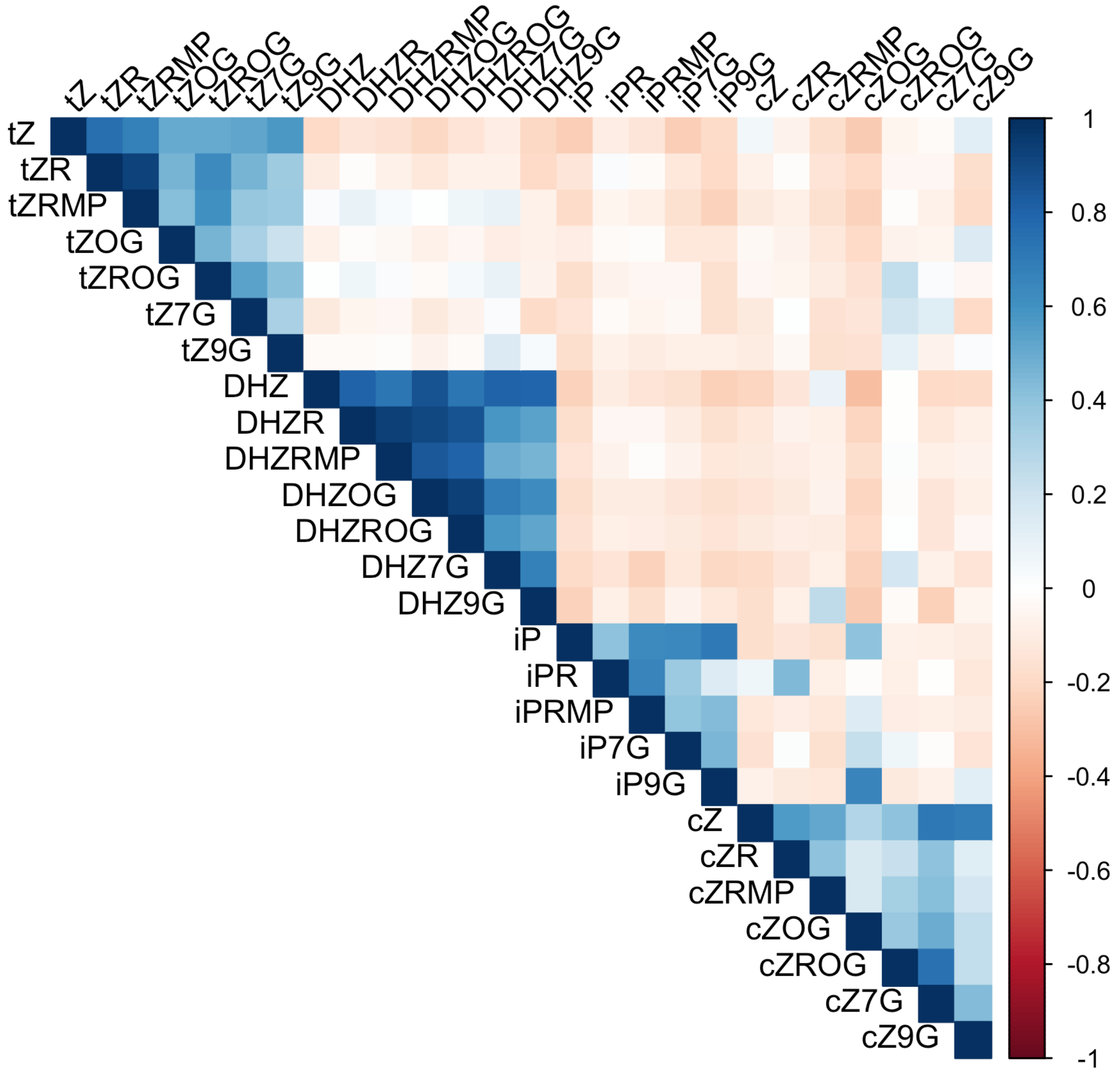

tZ
tZR
tZRMP
tZOG
tZROG
tZ7G
tZ9G
DHZ
DHZR
DHZRMP
DHZOG
DHZROG
DHZ7G
DHZ9G
iP
iPR
iPRMP
iP7G
iP9G
cZ
cZR
cZRMP
cZOG
cZROG
cZ7G
cZ9G
1
0.8
0.6
0.4
0.2
0
-0.2
-0.4
-0.6
-0.8
-1

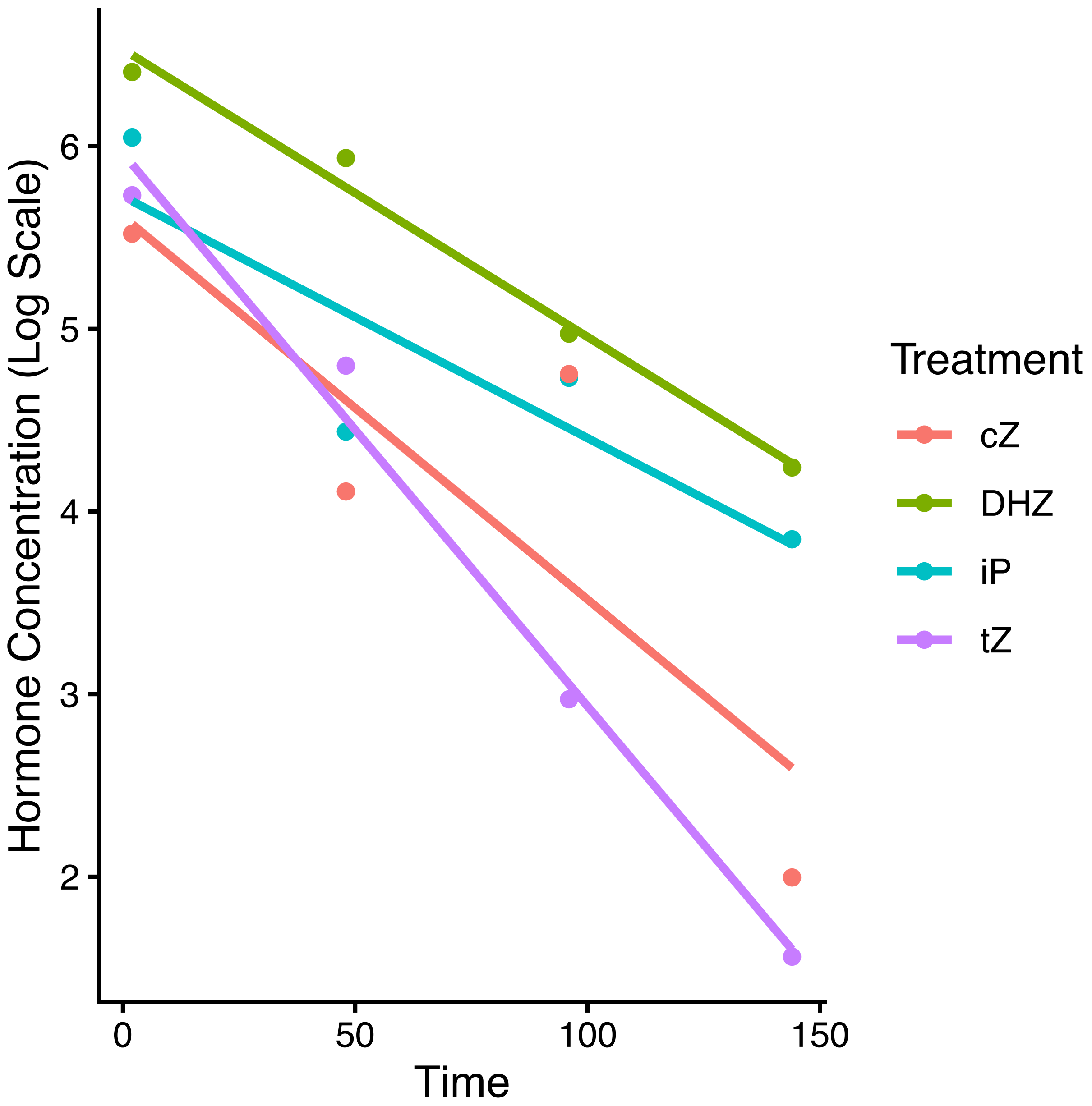

Hormone Concentration (Log Scale)
6
5
4
3
2
0
50
100
150
Time
Treatment
cZ
DHZ
iP
tZ

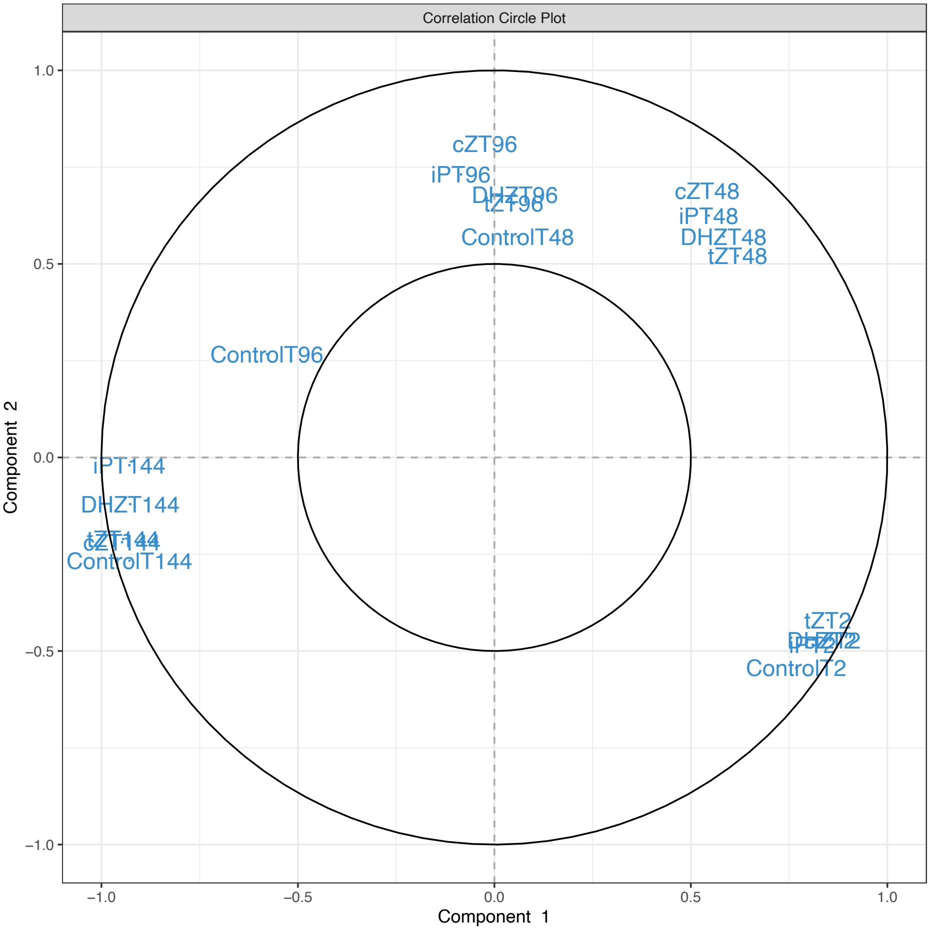
Correlation Circle Plot
cZT96
iPT96
DHZT96
tZT96
ControlT48
cZT48
iPT48
DHZT48
tZT48
ControlT96
iPT144
DHZT144
tZT144
cZT144
ControlT144
tZT2
ControlT2
1.0
0.5
0.0
−0.5
−1.0
−1.0
−0.5
0.0
0.5
1.0
Component 2
Component 1

A

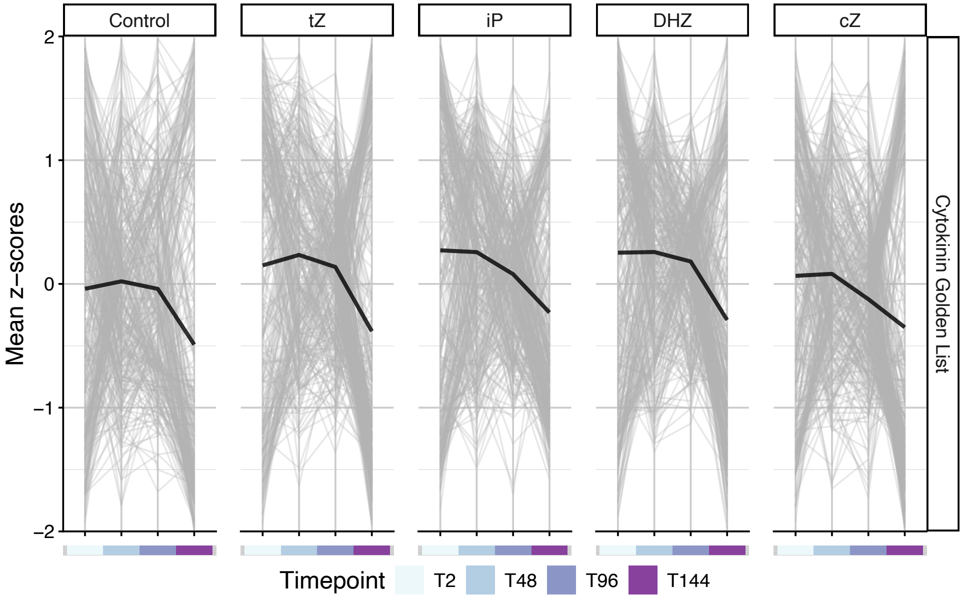

Control
tZ
iP
DHZ
cZ
2
1
0
−1
−2
Mean z−scores
Cytokinin Golden List
Timepoint
T2
T48
T96
T144


B

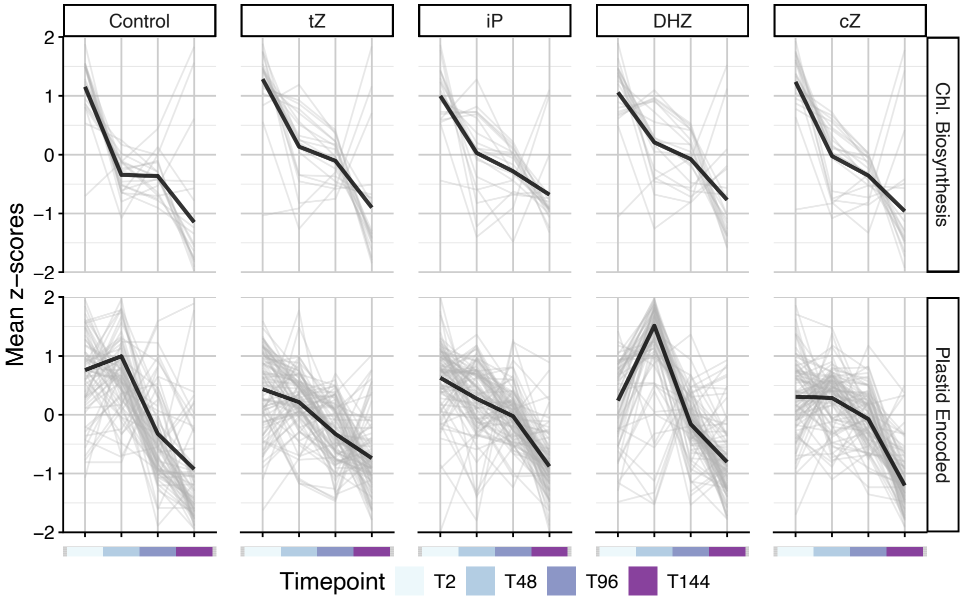

Control
tZ
iP
DHZ
cZ
2
1
0
−1
−2
Mean z−scores
Chl. Biosynthesis
Plastid Encoded
Timepoint
T2
T48
T96
T144

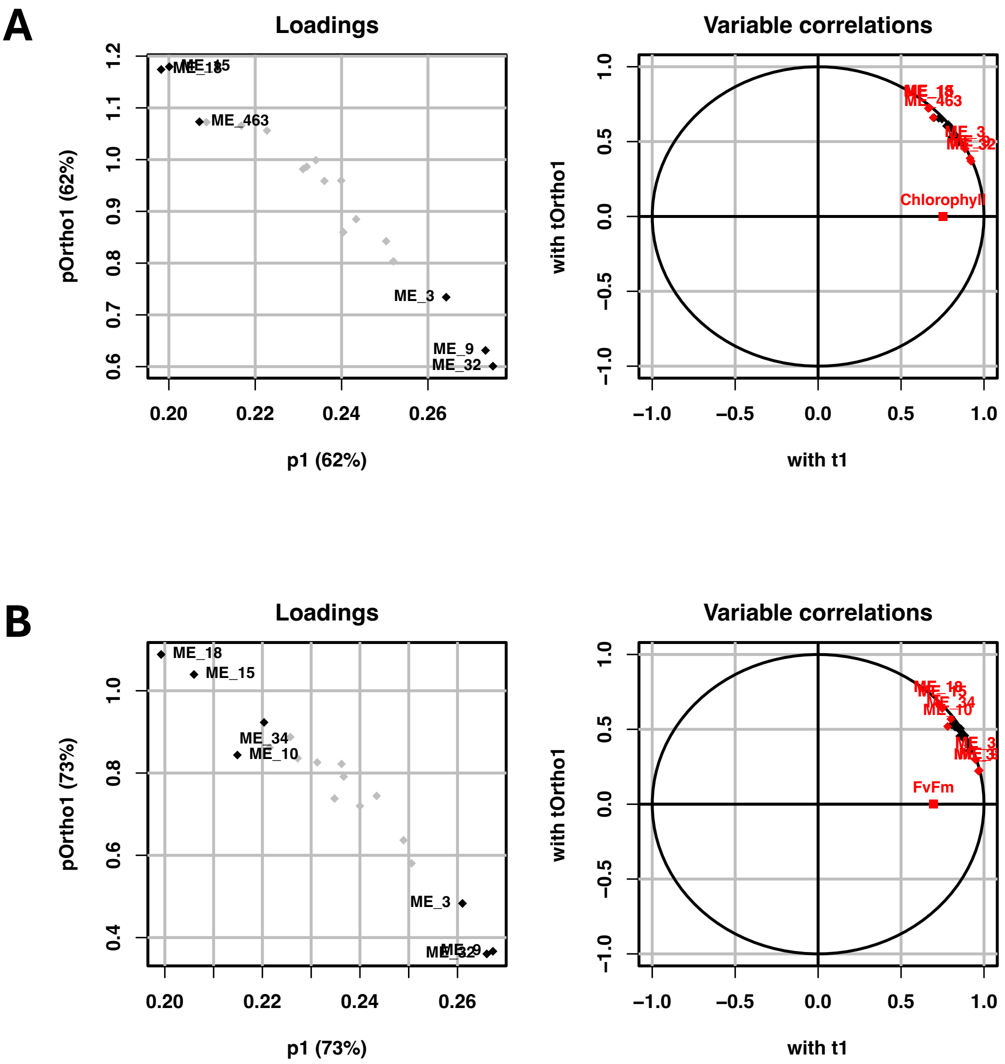
A
Loadings
pOrtho1 (62%)
p1 (62%)
ME_463
ME_3
ME_9
ME_32
Variable correlations
with tOrtho1
with t1
ME_463
Chlorophyll
B
Loadings
pOrtho1 (73%)
p1 (73%)
ME_18
ME_15
ME_34
ME_10
ME_3
Variable correlations
with tOrtho1
with t1
ME_10
FvFm

A

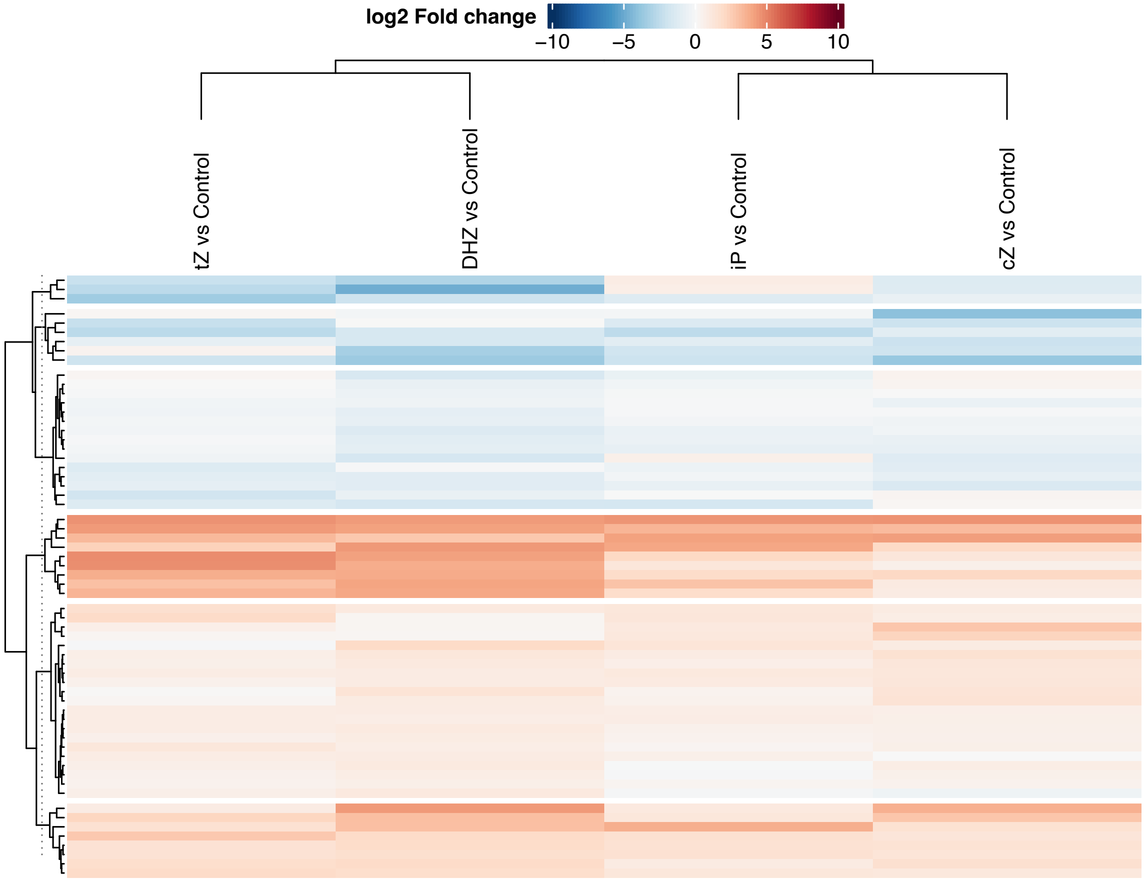

log2 Fold change
−10
−5
0
5
10
tZ vs Control
DHZ vs Control
iP vs Control
cZ vs Control
6
5
4
1
3
2


B

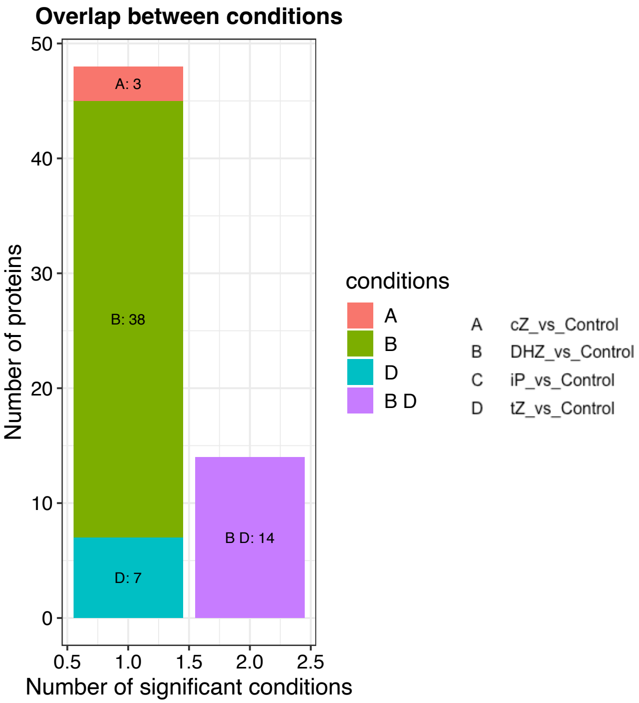

Overlap between conditions
A: 3
B: 38
D: 7
B D: 14
Number of proteins
Number of significant conditions
0
10
20
30
40
50
0.5
1.0
1.5
2.0
2.5
conditions
A
B
D
B D
A cZ_vs_Control
B DHZ_vs_Control
C iP_vs_Control
D tZ_vs_Control

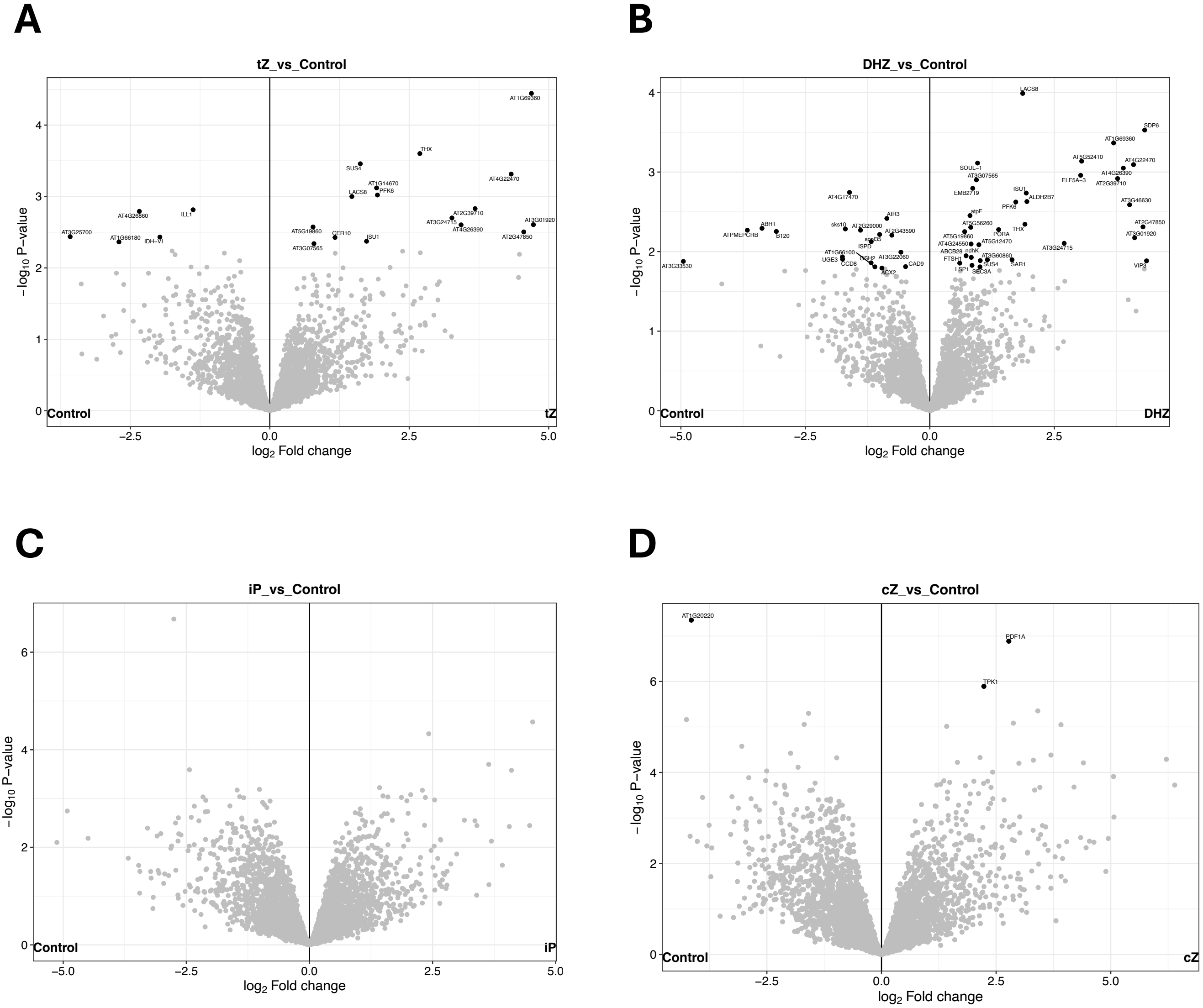

A
tZ_vs_Control
AT1G69360
THX
SUS4
AT4G22470
AT1G14670
LACS8
PFK6
AT4G26860
ILL1
AT2G39710
AT3G01920
AT3G24715
AT3G25700
AT5G19860
CER10
AT4G26390
AT1G66180
IDH-VI
ISU1
AT2G47850
AT3G07565
Control
tZ
−log10 P−value
log2 Fold change
B
DHZ_vs_Control
LACS8
SDP6
AT1G69360
AT5G52410
AT4G22470
SOUL-1
AT4G26390
AT3G07565
ELF5A-3
AT2G39710
EMB2719
AT4G17470
ISU1
ALDH2B7
AT3G46630
PFK6
AIR3
atpF
AT5G56260
AT2G47850
ABH1
sks10
AT2G29000
THX
AT3G01920
ATPMEPCRB
B120
AT2G43590
PORA
ISPD
AT5G19860
AT4G24550
AT5G12470
AT3G24715
AT1G66100
ABCB28
ndhK
AT3G22060
AT3G60860
UGE3
GSH2
FTSH1
SUS4
SAR1
VIP3
CCD8
CAD9
LSP1
SEC3A
AT3G33530
ACX2
Control
DHZ
−log10 P−value
log2 Fold change
C
iP_vs_Control
Control
iP
−log10 P−value
log2 Fold change
D
cZ_vs_Control
AT1G20220
PDF1A
TPK1
Control
cZ
−log10 P−value
log2 Fold change

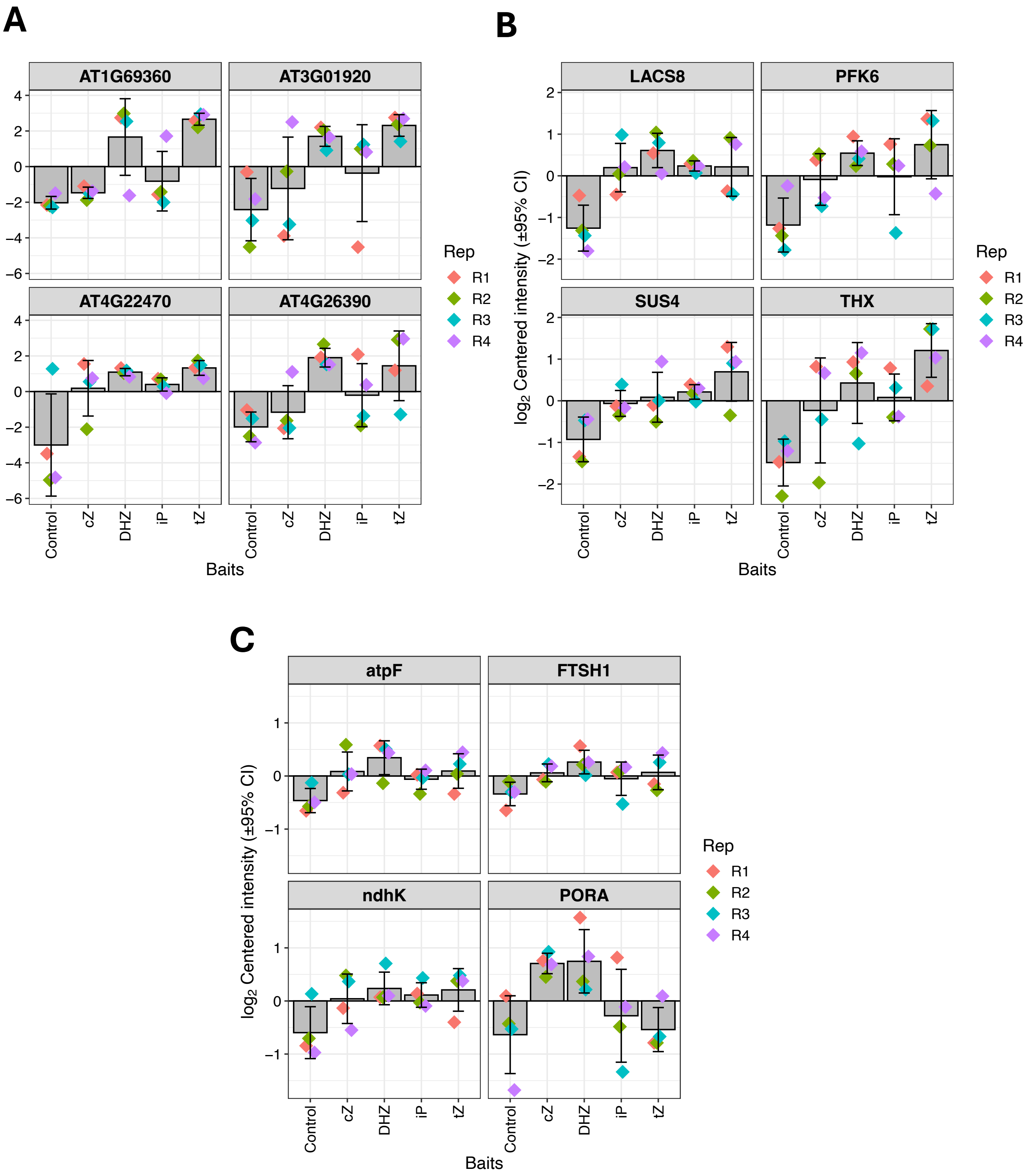

A
B
C
AT1G69360
AT3G01920
AT4G22470
AT4G26390
LACS8
PFK6
SUS4
THX
atpF
FTSH1
ndhK
PORA
log2 Centered intensity (±95% CI)
Baits
Control
cZ
DHZ
iP
tZ
Rep
R1
R2
R3
R4
